# Pathway Specific Unbinding Free Energy Profiles of Ritonavir Dissociation From HIV-1 Protease

**DOI:** 10.1101/2024.11.15.623815

**Authors:** Emily Vig, Jianan Sun, Chia-en A. Chang

## Abstract

Investigation of protein-drug recognition is key in understanding drug selectivity and binding affinity. In combination, binding/unbinding free energy landscape and intermolecular interactions can be used to understand the drug binding/unbinding mechanisms. This information is vital for the development of drugs with improved efficacy and explanation of mutation effects. This study investigated the dissociation processes of ritonavir unbinding from HIV protease (HIVp). Analyzing unbinding trajectories modeled by accelerated molecular dynamics (MD) simulations, three distinct pathways, Pathways A, B, and C, were characterized. Using reduced dimensionality strategy with the principal component analysis, we carried out short classical MD runs with explicit water to sample local fluctuation during ritonavir dissociation and applied the milestoning theory to construct unbinding free energy landscape. We found that each pathway showed similar values of binding free energy, albeit Pathway A accounts for over 50 percent of dissociation trajectories. Interesting, residue-residue correlation network analysis showed that in Pathway A, a broad correlation network outside the flap region governs protein motions during ritonavir unbinding which includes residues with reported mutation effects. However, the other two pathways showed limited correlation networks where no reported mutated residues were involved, explaining the favorability of Pathway A. Guided by the free energy profile, we investigated how and why of an energy barrier and minimum and demonstrated that hydrogen bonding governed the movement of the flap regions, directly impacting the calculated energy. Our study provided a new strategy to estimate ligand binding free energy and demonstrated the importance of the transient interactions during ligand-protein dissociation pathways in understanding drug unbinding.

## INTRODUCTION

Drug development relies heavily on a thorough understanding of how proteins and drugs interact (1). Knowledge of protein-drug interactions, both in bound and transient states, enables researchers to strategically develop drugs, thus improving their ability to selectively bind to proteins of interest (2,3,4). The dynamic nature of molecular interactions results in a complex environment in which many transient states occur over the course of drug binding and dissociation (5). These transient states arise within an energy landscape where different conformations correspond to different energy states. Various unbinding pathways often emerge as the system explores these energy minima and transition states during the unbinding process (6). Protein-drug interactions often differ between binding/unbinding pathways that contribute to their binding affinity; however, these intermediate processes are left unstudied by traditional approaches. Further, residues mutations, both in or outside the binding pocket, may induce structural and dynamic changes of proteins, which may have a substantial influence on protein-ligand interactions which impact drug potency, efficacy, and selectivity (7,8). Because of this, thoroughly investigating protein-drug interactions can provide explanations for drug resistance associated with protein mutations, both natural and induced.

Understanding the thermodynamic and kinetic properties of drug binding and unbinding is of fundamental importance for determining the efficacy of a drug (9). The dissociation rate constant, 1/k_off_, can be used directly to estimate the drug residence time (10). Residence time is a key indicator used in accessing drug efficacy, as drugs are only effective when bound to their respective target (11). Experiments measured thermodynamic and kinetic properties of protein-drug binding, such as rate constants (k_on_ and k_off_) and binding affinity (ΔG). While these important values are used to guide drug design, solely the measurements do not interpret and illustrate the complete picture of structure-kinetics and/or structure-affinity relationships in protein-ligand systems. To achieve a complete and thorough understanding of drug binding, computational methods can be used to illustrate and connect structural interactions with thermodynamics, kinetics, and mechanisms of molecular processes. With these measured properties we can strategically plan a drug development process to improve both specificity and efficiency. This is of extreme interest in current drug development in which druggable targets have been identified, including human immunodeficiency virus (HIV).

HIV is a potentially life-threatening virus that attacks the human immune system and, when left untreated, results in acquired immunodeficiency syndrome (AIDS). A few viral proteins have been utilized for AIDS treatments, such as HIV-1 protease (HIVp) (12). However, as the viral proteins constantly mutate, the development of drugs to kill HIV has proved difficult. HIVp is a symmetrical homodimeric protein responsible for the cleavage of premature viral proteins (13). This cleavage is responsible for the formation of mature, infectious HIV particles (14). HIVp serves as a prominent model system in scientific research and drug development (15)(**Figure 1A**). The well-defined three-dimensional structure of HIVp allows insight into its molecular functions, facilitating the modeling and optimization of inhibitors in rational drug design (16). Protease inhibitors, a class of antiretroviral drugs, work by obstructing HIVp and inhibiting its function. One unique feature of HIVp is the well-documented characteristic flap movement, opening and closing, which serves as a gate to govern natural substrate and drug binding/unbinding (17,18). Understanding the flap movement is crucial for designing effective inhibitors (19). Many HIVp inhibitors work by binding to the enzyme’s active site and stabilizing the closed state (20–22). This closed state prevents the flaps from opening, thus inhibiting the enzyme’s catalytic activity. This prevents the formation of infectious viral segments thereby limiting the progression of HIV infection. Studying the dynamics of flap movement and different drug binding/unbinding pathways with the flap dynamics provides insights into the molecular mechanisms of HIVp inhibitor binding and aids in better developing antiretroviral drugs incorporating this dynamic process (23). The mechanism of action of these HIVp inhibitors serves as a model for understanding enzyme inhibition, contributing to broader knowledge applicable to other enzymatic systems.

**Figure 1.**
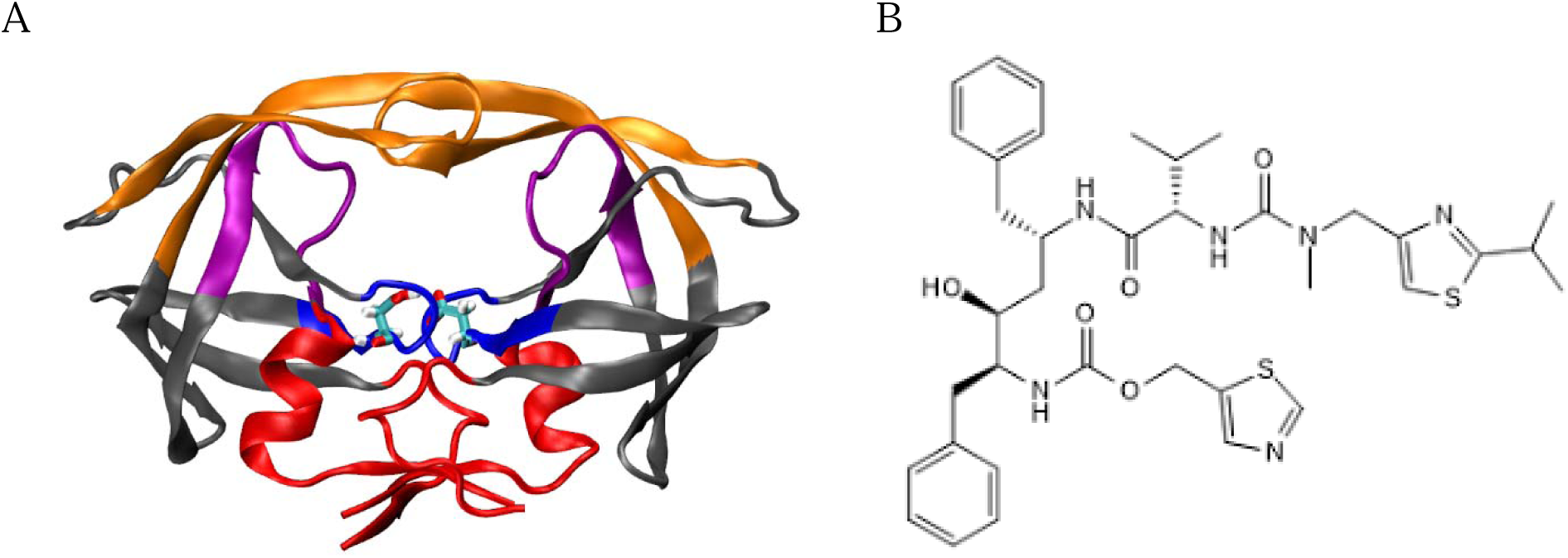
HIVp and Ritonavir Structures. **(A)** HIVp structure with color-coded regions: flap (orange: residues 42–60 and 141–159), flap tips (orange: residues 49–52 and 148–151), loop (pink: residues 75–85 and 174–184), catalytic triads (blue: residues 23–29 and 122–128) and dimer interface (red: residues 1–9, 86–99, 100–108 and 185–198). Catalytic residues, Asp 25 and Asp 124, are shown in licorice. **(B)** Two-dimensional structure of HIVp inhibitor ritonavir

Ritonavir is a common HIVp inhibitor that is used in the treatment of HIV (**Figure 1B**)(24). Ritonavir non-covalently binds to HIVp and stabilizes the closed state. Experimental techniques (i.e. X-ray Crystallography) determine 3-dimension structures that allow observation of ritonavir bound to HIVp (25). However, crystal structures lack movement and do not give insights into all the interactions that occur during the unbinding of ritonavir from HIVp. This is particularly important when flap mobility is key in drug-unbinding. While Nuclear Magnetic Resonance (NMR) spectroscopy can provide quantitative information of protein dynamics, NMR has limitations in revealing all atomistic details during drug binding/unbinding. Understanding these dynamic interactions is crucial, as HIVp mutations, which affect flap mobility, have been shown to increase resistance to ritonavir (26). Insights into ritonavir’s dynamic interactions with HIVp may help to explain this resistance.

Molecular dynamics (MD) simulations model movements of protein-drug complexes over time, providing insights into the interactions and structural changes associated with drug association/dissociation (27). MD allows for the visualization of unbinding pathways and can handle large, complex systems, making it ideal for understanding dynamic processes and resistance mechanisms. In conventional molecular dynamics (cMD) simulations, the time it takes for a drug to dissociate from a protein can vary greatly. For strong binders, dissociation might take milliseconds to hours. As cMD typically only simulates nanoseconds to microseconds of real-time dynamics, observing spontaneous dissociation within this period is highly unlikely, unless additional techniques are used to speed up the process. Enhanced sampling techniques have been widely used to investigate ligand binding kinetics (28). For example, using accelerated molecular dynamics (aMD) has allowed efficient conformational sampling of HIVp by reducing energy barriers which resultingly increases the occurrence of rare events (29,30). In combination with a reseeding approach, aMD can be used to collectively study rare dissociation events in great numbers. Earlier studies using this approach have shown that ritonavir dissociates from the binding pocket via three distinct pathways (31). Though past simulations have shown the presence of multiple pathways, additional analysis and constructing ligand unbinding free energy landscape further provide quantitative data to understand protein-drug recognition and the probability differences between each pathway. Popular analyses such as Molecular Mechanics Poisson-Boltzmann Surface Area (MM-PBSA) provide insights into molecular interaction energy and solvation effects (32). However, changes of entropy and free energy during ligand binding/unbinding, the vital assessment of molecular flexibility, are difficult to calculate via traditional techniques. As HIVp is known to function largely through flap movements, accurately describe protein flexibility is crucial in fully understanding ligand binding and protein function.

A key metric for understanding the thermodynamics of the complex is the free energy profile, also termed Potential of Mean Force (PMF). Here PMF refers to the free energy of a system as a function of a defined coordinate. It is commonly seen for ligands to move back and forth in the binding pocket, as is the case with ritonavir **(Figure S1)**. Therefore, providing a defined coordinate which can accurately describe the molecular movement during drug unbinding is not straight forward. For example, use of simple center of mass distance or root mean square deviation (RMSD) of ligand position may not be adequate to depict ligand binding or unbinding. As a result, reduced dimensionality using principal component (PC) analysis has been utilized to better define a ligand unbinding coordinate (33).

In this study, we calculated the free energy profile for ritonavir unbinding from HIVp. We investigated 3 major unbinding pathways and used the Binding Kinetics Toolkit (BKiT) package to define the ritonavir unbinding coordinate and construct PMF using many short cMD and the milestoning theory (34,35). The exact drug-protein interactions from local energy minima are of extreme interest as they would affect drug-binding kinetics, even though the absolute binding free energy is the same despite the unique unbinding pathways. Protein-drug conformations existing in minima stabilize the HIVp-ritonavir complex. By connecting local free energy minima and barriers to key unbinding events, we can verify the free energy profiles and explain how hydrogen binding in flap regions impacts free energy. Energy wells are directly attributed to hydrogen bonding between ritonavir and HIVp stabilizing the bound complex. Further, sidechain correlation studies provide insight into total protein movement and cohesiveness unique to each of the three pathways. Understanding protein movement and ligand-protein interactions at transient states can help us optimize drugs with desired kinetic properties (36,37) as well as provide insight into the cause of mutation-related drug resistance.

## METHODS

### Molecular systems

Ritonavir-HIVp dissociation trajectories sampled by aMD were taken from published work that used Amber14SB for protein and GAFF for ligand (31). The initial frame of aMD was taken from 100-ns conventional MD (cMD) runs, which used the protein data bank entry 1HXW as the initial bound state, with a single protonation state applied for Asp25. AMD allows for the sampling of rare dissociation events by adding a continuous non-negative bias or “boost” potential. Both dihedral and total potential energy boosts were applied. Boost factors were calculated based on 100-ns bound state ritonavir-HIVp cMD simulations. Re-seeding was used to generate multiple MD simulation production runs from one initial conformation. AMD, when combined with re-seeding, results in the capturing of many ritonavir unbinding events. In order to better sample these unbinding events, we selected frames from the aMD dissociation trajectories as initial frames for short MD runs. Initial frames for short MD runs were taken every 100ps for the majority of the simulation. As ligand leaves the binding pocket, we decrease the interval of initial frame selection to once every 20ps to better cover the PC space. Initial frames varied between pathways to account for differences in simulation length. The breakdown of initial frames is detailed in **Table S1**. We ran 10 replicas of short MD runs for each initial frame. Explicit solvent was used. All short MD runs were 100 ps long unbiased conventional MD with TIP3P water model, Amber14SB for protein, and GAFF for ritonavir at 298K. Frames in short MD runs were saved every 100 Fs. By using many short MD runs we are able to (1) sample transitions between unbinding indexes and (2) validate proposed dissociation pathways.

### Constructing the unbinding free energy profile

#### Defining Unbinding Coordinates

To project high dimensional trajectories into low dimensional spaces, cartesian coordinates are taken from aMD and stored in a matrix consisting of the x,y,z coordinates of each alpha carbon of HIVp and each heavy atom of ritonavir. This matrix is then projected onto the Principal Component (PC) space. Only the first two PCs (PC1 and PC2) are generated. PC space encapsulates the key motions of ritonavir into a smaller subset of principal components. This results in a series of positions that represent the primary motions ritonavir undergoes during dissociation from HIVp. A score plot is made of PC1 vs PC2 and a smooth path is generated by taking a continual average of frames before and after 100 steps of each frame. Loops are then removed from the smoothed path and milestones are placed with a width of 4 angstroms. Each milestone represents an unbinding index, a key position of ritonavir during the dissociation from HIVp.

#### Defining Representative Frames

Representative clusters for unbinding index are then defined as projected frames that fall within 1.5 angstroms of a milestone. For any given unbinding index, we keep at most 10 configurations that fall the closest to the milestone. These frames are saved for validation of unbinding trends associated with each milestone. A trajectory can then be created using one representative frame for each milestone. For milestones with multiple frames per representative cluster, we selected the frame nearest to the average in PC1/PC2 space as the representative frame. For milestones with no representative clusters, we use a representative frame from the neighboring milestone.

#### Construction of the Transition Matrix

Transitions between the unbinding indexes are computed by analyzing system behavior in numerous short MDs. We provide two methods of constructing transition matrix, PC based or ligand RMSD based. For the PC based method, unbinding indexes are found as previously described. The system movement in each short MD is examined and judged to see if it crossed any two consecutive unbinding indexes. If a transition occurs, two neighbor unbinding indexes and frames are recorded to compute the time needed in such transition. BKiT counts numerous transitions and computes the average lifetime (τ*_i_*) and its standard deviation (σ_τ,i_) of each transition. This data then is used to construct a transition matrix, which quantifies the probability of transition in or out of each unbinding index. The ligand RMSD based method uses the same protocol for constructing the transition matrix, instead using ligand RMSD as initial unbinding indexes.

#### Constructing the unbinding free energy profile

The free energy profile, also termed potential of mean force (PMF), provides quantitative free energy values using the unbinding index as a reaction coordinate. This allows us to examine energy trends relative to each stage of ritonavir’s dissociation. Free energy of each unbinding index can be calculated by: F = -K_B_T ln (q_i_ t_i_) where q_i_ is the stationary flux of each unbinding index (i) and t_i_ is the averaged lifetime of unbinding index (i). Stationary flux quantifies the number of transitions into any given unbinding index per unit time, as computed from the transition matrix. Bkit computes the stationary flux by solving the eigenvalues of the transition kernel. Eigenvalues equal to 1, the highest possible value, represent the stationary flux. The mean first passage time (MFPT) of index n can be estimated by: MFPT = ∑ q_i_ t_i_ / q_n_, where residence time is estimated when the index n shows that the ligand already unbinds from the pocket.

### Examining Ligand-Protein Interactions and Stability Factors

#### Hydrogen Bond Analysis

The unbinding free energy is largely determined by interactions within the protein. For Instance, each hydrogen bond between the ligand and protein may contribute up to a few kcal/mol to interaction energy (38). To hydrolyze the peptide bond in premature protein, Asp25/124 establishes hydrogen bonding with the peptide backbone in the substrate, whereas Ile50/149 stabilizes the substrate with a water bridge (39). To better understand which residues may form H-bonds with ritonavir during ligand dissociation, we analyzed and plotted H-bond versus time by using the CPPTRAJ program (40). The criteria for H-bonding are (1) the distance between donor D and acceptor A < 3 Å, and (2) the D-H-A angle, where H is the shared hydrogen, at least 150◦. Hydrogen bonds were also collected using the VMD hydrogen bond plug-in, following the same parameters.

#### Pairwise Sidechain Correlation

Sidechain correlation values were measured using T-Analyst (41). The Pearson correlation formula was used to compute pairwise correlations. To ensure accuracy and avoid computational errors arising from discontinuity margins (±180° or 360°/0°), we transformed the side-chain rotation angles into Cartesian coordinates using equations (1)-(3).

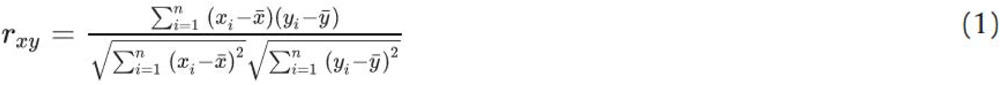

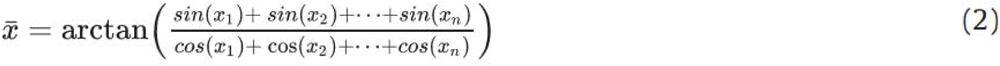

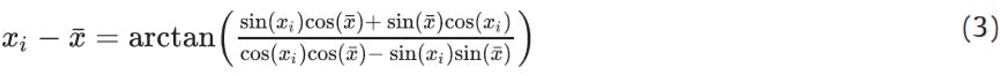

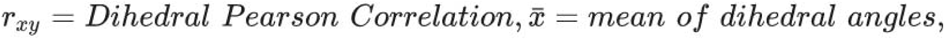

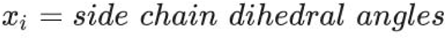

Residues that shared a correlation value above 0.50 were labeled positively correlated, with negatively correlated values falling below -0.50. A positive correlation between two sidechains suggests that they rotate similarly during MD simulation. As tail regions show high movement, there is increased odds of capturing correlation by chance. For this reason, correlation pairs including tail regions have been negated.

## RESULTS AND DISCUSSION

This section begins by reporting ritonavir unbinding free energy landscape for distinct unbinding pathways sampled by aMD runs (31). Pathway A describes unbinding between the flap region and the loop region. Pathway B describes surface-level diffusion through the flap region. Pathway C describes surface-level diffusion through the interface region. Of these studies, Pathway A accounted for over 50 percent of dissociation trajectories. We showed that ritonavir’s navigation through HIVp in each pathway and drug-protein interactions associated with each pathway differ greatly. We used two unbinding coordinates to present unbinding index for constructing PMF. The investigation of residue-residue correlation and the implications of mutation effects are also discussed.

### Computed unbinding free energy landscape

#### Using PC space to compute free energy landscape and residence time

Using a coordinate to precisely describe ligand unbinding from the protein pocket is crucial in PMF calculations. Only when the unbinding coordinate can be reasonably described, which include key movements of protein sidechains and ligand functional groups, the free energy barriers and minima shown in the computed free energy landscape can accurately associate with ligand dissociation processes and explain binding kinetics. As demonstrated in **Figure S3**, we first performed principal component analysis (PCA) and project the Cartesian coordinate representation with high-dimensional ligand-protein motions (3N-6 degrees of freedom, where N is the number of atoms) to only 2-dimention (2D) space using the first 2 PC modes (degrees of freedom). We used the BKiT package to assign milestones in the 2D-PC space, and construct PMF using various short 100ps classical MD runs.

As illustrated in **Figure 2**, the calculations revealed rugged ritonavir unbinding free energy landscape, where the free energy barriers and minima shown in the landscape associate to important molecular movements, as discussed in the next subsection. Unbinding free energies and residence time of ritonavir-HIVp were also be approximated from the PMF (**Table 1**, **left column)**. PC modes represent fundamental motions during ritonavir dissociation, where Pathways A and B share similar unbinding free energy values, -9.4 kcal/mol and -9.5 kcal/mol, respectively. Pathway C is less energetically favorable, with an unbinding free energy of -6.1 kcal/mol. Pathways A and B demonstrate a gradual increase in energy as the unbinding progresses (**Figures 2A and 2B**), while Pathway C shows a large and shallow free energy well during ligand unbinding (**Figure 2C**).

**Figure 2.**
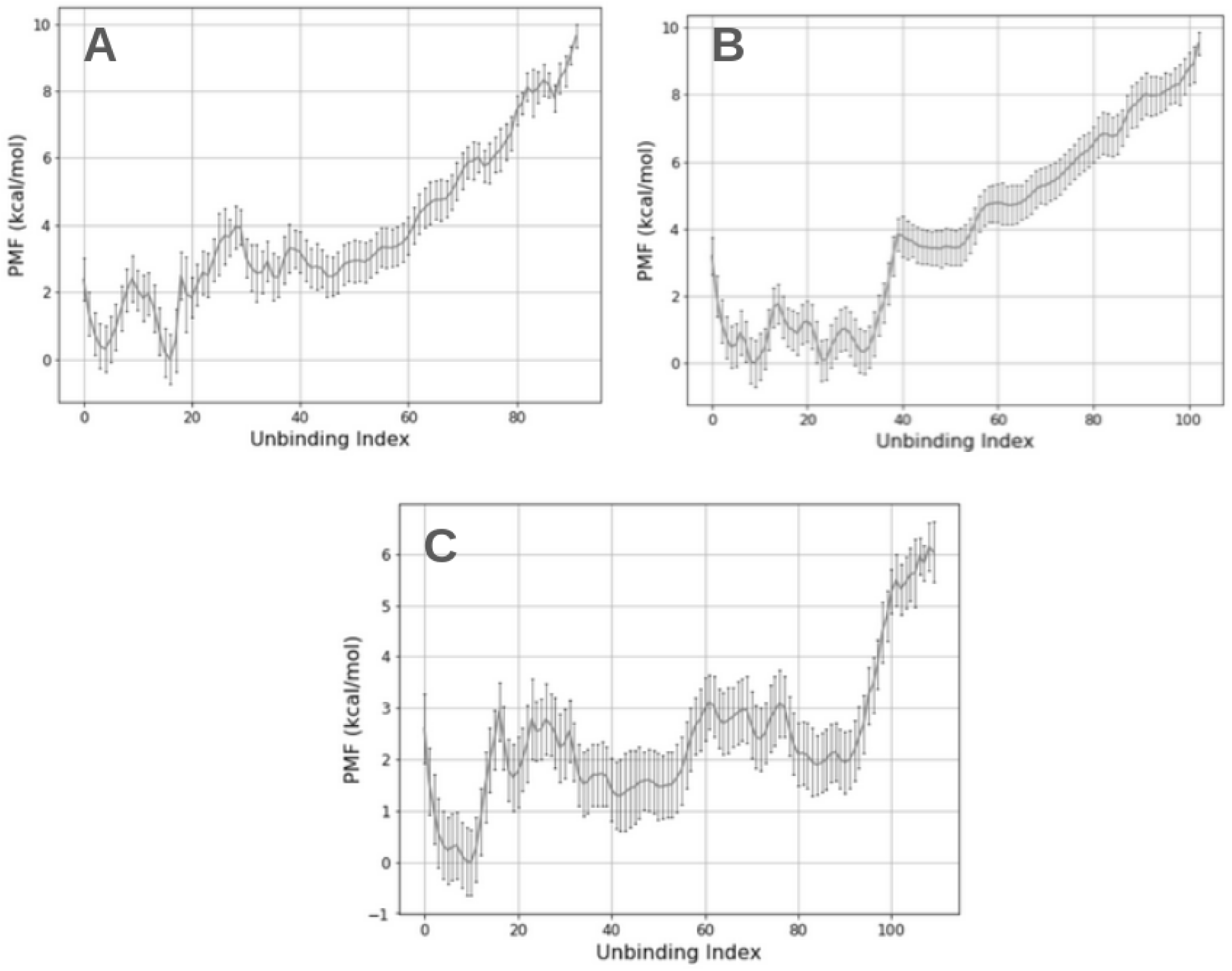
Pathway specific free energy landscape of ritonavir unbinding from HIVp. Free energy landscape of pathway A (A), pathway B (B), and pathway C (C) computed using unbinding coordinates defined from the PC space, labeled as PC coordinate in Table 1. Pathways A and B demonstrate a progressive increase in energy. Pathway C exhibits an initial jump in energy, followed by a large plateau, before a final energy jump upon unbinding. The absolute binding free energy ΔG estimated from the plots is systematically underestimated, compared with experimental results. However, the free energy plots allow for relative energy studies of local minima and maxima.

**Table 1.** Calculated and experimental unbinding free energy values and pathway occurrence. Pathway-specific binding free energy (kcal/mol) and residence time computed with coordinates assigned using the PC space (labeled PC Coordinate) and ligand RMSD (labeled RMSD Coordinate) in comparison to experimental values. Percentages show frequency of each pathway out of a total of 20 successful ligand dissociation runs modeled by aMD simulations (31). Percentages do not add up to 100% due to negation of undefined pathways.

|  | *PC Coordinate* |  | *RMSD Coordinate* |  | *Experimental* |  | *aMD* |
| --- | --- | --- | --- | --- | --- | --- | --- |
| | Binding Free Energy ($\Delta G$) | Residence Time | Binding Free Energy ($\Delta G$) | Residence Time | Binding Free Energy ($\Delta G$) | Residence Time | Occurrence |
| *Pathway A* | -9.4 | 483 μs | -14.1 | 2.5 s | -13.7 | 463 s | 55% |
| *Pathway B* | -9.5 | 2 ms | -15.4 | 41.6 s | -13.7 | 463 s | 20% |
| *Pathway C* | -6.1 | 4.5 μs | -14.9 | 29.6 s | -13.7 | 463 s | 15% |

#### Using ligand RMSD to compute free energy landscape and residence time

While the PMF using the 2D PC space to present the unbinding coordinate provides valuable insights into HIVp and ritonavir movement during ritonavir unbinding, the absolute binding free energies from all 3 pathways were underestimated, compared with the experimental binding free energy ∼ -13.7 kcal/mol. The respective residence times also differed from the experimental value of 463 s. As detailed in a published work, the differences between the computed and experimental values mainly came from skipping data points when counting transitions between milestone indexes presented by straight lines in the 2D space **(Figure S7)** (42). In addition, missing coverage of molecular motions using the first 2 PC modes. The combined coverage of the first 2 PC modes for ritonavir unbinding from HIVp under pathway A was 56%, where 44% of the motions were not considered in the 2D space. Pathways B and C had 69.8% and 60.9% coverage, respectively. Therefore, to avoid skipping data point when using a line in the 2D space, we used ligand RMSD to present ritonavir unbinding with 0.1Å interval to present milestones. This 1-dimension (1D) milestone index can ensure that every transition between two adjacent milestones is counted, and no data point is skipped. The computed binding free energy with the 1D milestone index under pathway A, B, and C are -14.1 kcal/mol, -15.4 kcal/mol, and -14.9 kcal/mol, respectively (**Table 1**, **middle column)**, which are in good agreement with experimental measurement. The computed drug residence time for ritonavir dissociation from HIV protease under pathway A, B, and C are 2.5 s, 41.6 s, and 29.6 s, respectively, which are in the ballpark of experimental residence time, ∼ 463 s. While use of 1D to present milestones can accurately reproduce experimental binding free energy and residence time, using ligand RMSD cannot accurately describe HIVp and ritonavir motions during ligand dissociation (**Figure S1, Figure S2**). Nevertheless, this correction validates that the error sources came from use of milestones presented by lines in the 2D PC space, which inevitably skip transition counts.

#### Thermodynamic investigation of free energy landscape

By examining the free energy landscape, we are able to illustrate the relative stability of different HIVp-ritonavir conformations and provide insights into how and why the transitions between the free energy minima and barriers. Here we aim to examine conformations associated with these energy minima and barriers to learn the governing key interactions that occur during ritonavir dissociation. We showed that these interactions are different between dissociation pathways, which also influence the overall movement of HIVp. Further, by being able to connect energy data to existing interactions, we validate both the unbinding pathway as well as the free energy landscape.

#### Free energy landscape under pathway A

The unbinding free energy landscape identifies three major energy wells and three major energy barriers over the course of ritonavir’s dissociation from HIVp via pathway A (**Figure 3**). The computed free energy landscape correctly captured the interactions of the crystal-bound state at energy minimum A (**Figure 4A**). Upon visual analysis, it was concluded that the computed free energy landscape can correctly capture the crystal bound state. By visualizing ritonavir-HIVp interactions at energy minima A, we found hydrogen bonds between ritonavir and Asp25 and Asp 29, in addition to pi-stacking between ritonavir and Arg107. As the system progressed, interactions between ritonavir and HIVp were broken, giving us an energy barrier B (**Figure 4B**). As ritonavir approaches Thr 80, a strong hydrogen bond of 1.8 angstrom between the hydroxyl groups of ritonavir and Thr 80 was formed, which further attracted ritonavir to loop region of Thr 80 (**Figure 4C**). More interactions were followed by such hydrogen bonds. Gly 150 and Ile 149 formed hydrogen bonds with ritonavir as well. Non-polar attraction between Ile 146 and ritonavir was also observed. Flap regions of HIVp opened, exposed ritonavir to solvent, and reduced contact with ritonavir, giving us an energy barrier D (**Figure 4**). Non-polar attraction was the major interaction between ritonavir and HIVp. As ritonavir keeps moving to left side of HIVp, flap region on the right side closed, resulting in a slight lower energy barrier compared to barrier D (**Figure 4E**). Once ritonavir moved to the gap between flap and loop region, a local energy minimum was observed at energy well F (**Figure 4F**). As ritonavir kept moving along unbinding pathway A, it temporarily formed hydrogen bond with Lys 154, resulting in a tiny energy minimum at G (Figure 4G), prior to ritonavir’s complete dissociation.

**Figure 3.**
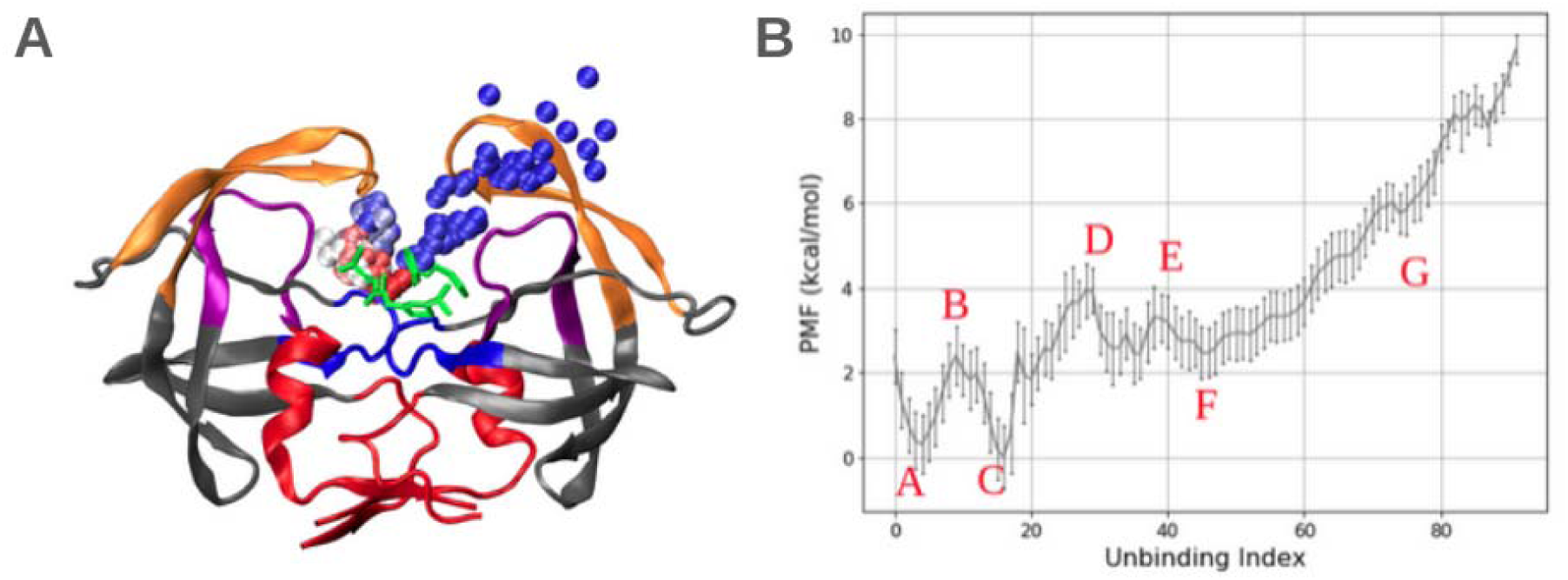
Free energy landscape of ritonavir unbinding from HIVp under pathway A. **(A)** Ritonavir dissociation under pathway A. HIVp colored as in **Figure 1**. Each bead represents a position of center of mass of ritonavir with 1-ns interval taken from aMD dissociation, with total simulation time of 248.7 ns. The HIVp conformation is taken from the final frame of the dissociation trajectory. Color beads present center of mass of ritonavir taken from frames in the beginning (red), middle (white) and near the end (blue) of the trajectory. Ritonavir’s initial position is green licorice. **(B)** Free energy landscape of ritonavir under pathway A with labeled maxima and minima. Ritonavir Major energy barriers (B,D,E) and local minimum (A,C,F,G) labeled in red, and the conformations are detailed in **Figure 4**.

**Figure 4.**
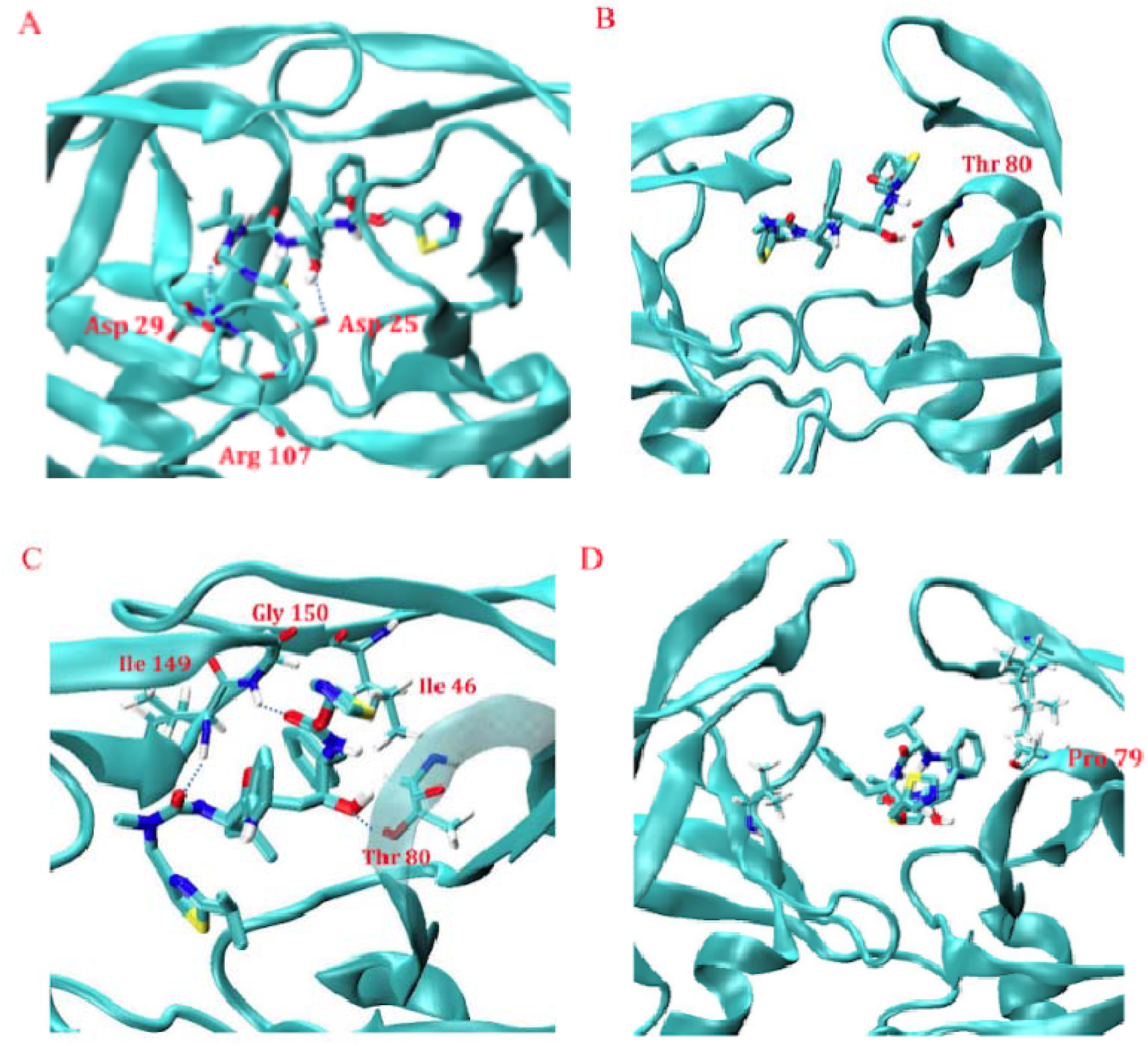
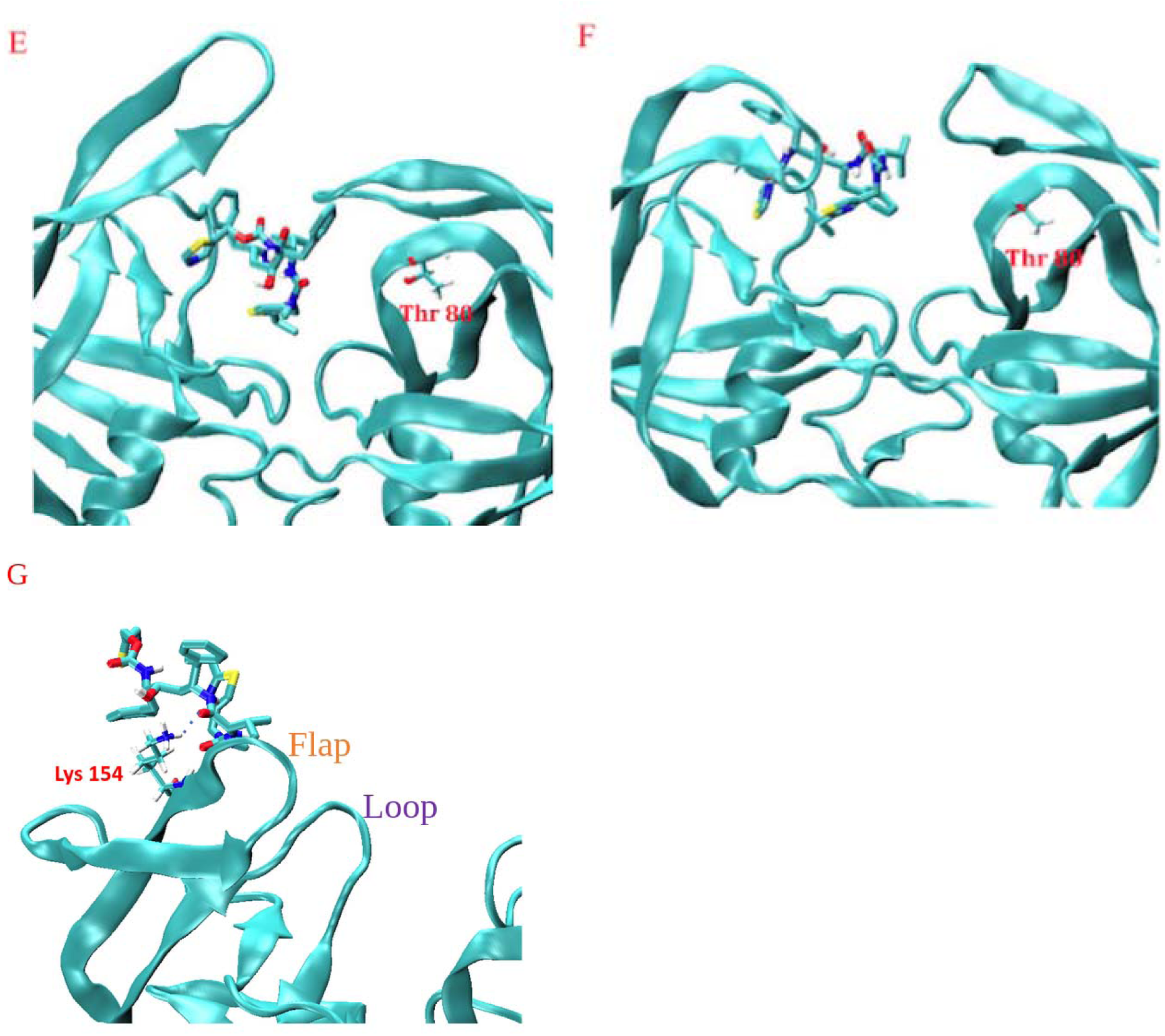
Progression of molecular interactions of Ritonavir and HIVp during dissociation via pathway. **A.** A-G show local energy minima or barriers indicated in **Figure 3**. **(A, C, G)**. Local free energy minima are explained via bound-state stabilization by Hydrogen bonds between ritonavir and HIVp. **(B, D, E)** Free energy barriers are seen when these Hydrogen bonds are broken, and ritonavir is positioned that allows high solvent exposure. **(F)** A free energy minimum reveals a favorable ritonavir positioning between the flap and loop regions.

#### Free energy landscape under pathway B

We further constructed the free energy landscape of ritonavir unbinding from HIVp under pathway B (**Figure 5**). Once again, the computed free energy landscape correctly captured the interactions of the crystal bound state at energy minimum A (**Figure 6A**). These hydrogen bonds were broken at energy barrier B (**Figure 6B**). Since pathway B involves ligand diffusion on flap region, minimal interactions between ritonavir and loop region were observed. At energy minimum C, ritonavir formed hydrogen bond with Asp 128, Gly 51 and Gly 148 (**Figure 6C**). Energy well D was observed with hydrogen bond between ritonavir and Asp 128, Ile 149 (**Figure 6D**). As flap regions opened widely and exposed ritonavir to solvent, reduced contact between ritonavir and HIVp and broken hydrogen bonds, a steep energy barrier showed up from D to E, where minimal contact between ritonavir and HIVp presented at energy barrier E (**Figure 6E**). As ritonavir moved along the flap region, it encountered Lys 142 and formed a transient hydrogen bond, giving a small energy well F (**Figure 6F**).

**Figure 5.**
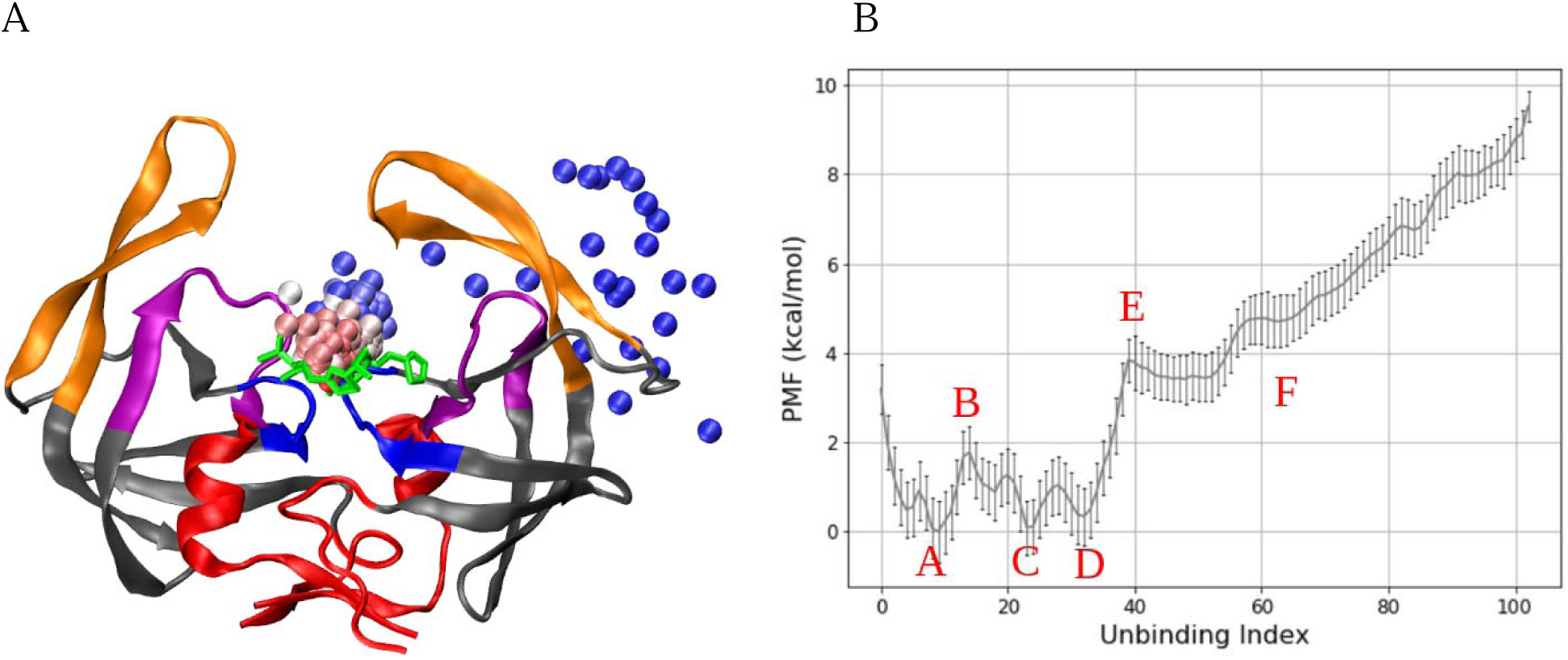
Free energy landscape of ritonavir unbinding from HIVp under pathway B. **(A)** Ritonavir dissociation under pathway B. HIVp colored as in Figure 1. Each bead represents a position of center of mass of ritonavir with 1-ns interval taken from aMD dissociation, with total simulation time of 304.9 ns. The HIVp conformation is taken from the final frame of the dissociation trajectory. Color beads present center of mass of ritonavir taken from frames in the beginning (red), middle (white) and near the end (blue) of the trajectory. Ritonavir’s initial position is green licorice. **(B)** Free energy landscape of ritonavir under pathway B with labeled maxima and minima. Major energy barriers (B,E) and local minimum (A,C,D,F) labeled in red, and the conformations are detailed in **Figure 6**.

**Figure 6.**
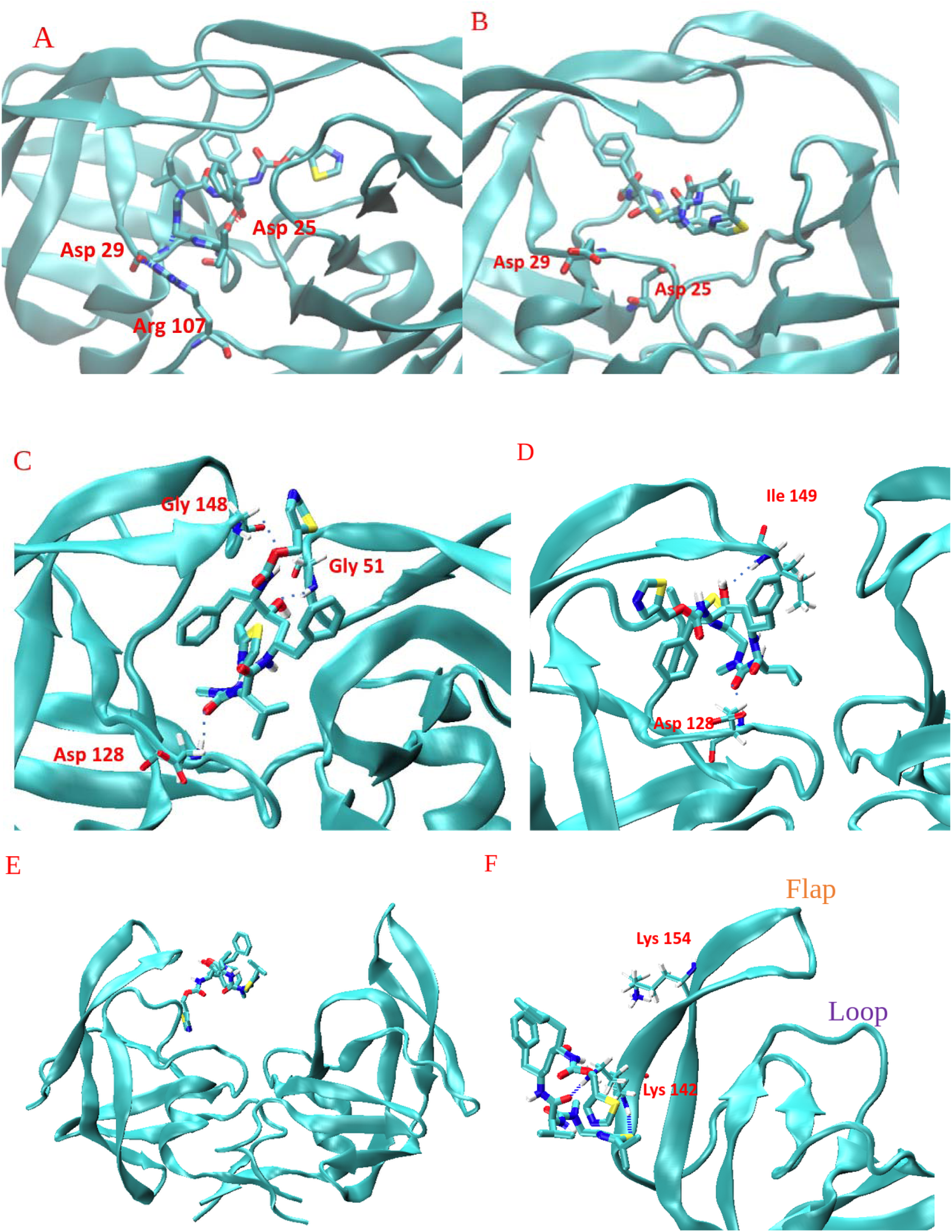
Progression of molecular interactions of Ritonavir and HIVp during dissociation via pathway. **B.** Local energy minima or barriers are indicated in **Figure 5**. **(A, C, D, F)** Local energy energy minima are explained via bound-state stabilization by Hydrogen bonds between ritonavir and HIVp. **(B,E)** Local free energy barrier has appeared when these bonds are broken, and ritonavir is positioned that allows high solvent exposure.

#### Free energy landscape under pathway C

We further examine the free energy landscape of ritonavir unbinding from HIVp under pathway C (**Figure 7**). The computed free energy landscape correctly captured the interactions of the co-crystal bound state of ritonavir-HIVp (**Figure 8A**) and the breaking of these hydrogen bonds at energy barrier B (**Figure 8B**). At local energy minimum C, ritonavir was mostly attached to the flap region with hydrogen bonds to Ile 50, Ile 149, and Gly 151 (**Figure 8C**). More interactions between ritonavir and interface region of HIVp were observed. At the energy well D. Ritonavir left the flap region and started contacting Arg 8 with two hydrogen bonds, plus hydrogen bonds with Asp 129 (**Figure 8D**). As ritonavir kept moving towards the interface region, the attraction between ritonavir and Arg 8 grew strong. Three hydrogen bonds were observed between ritonavir and Arg 8 at energy well E (**Figure 8E**).

**Figure 7.**
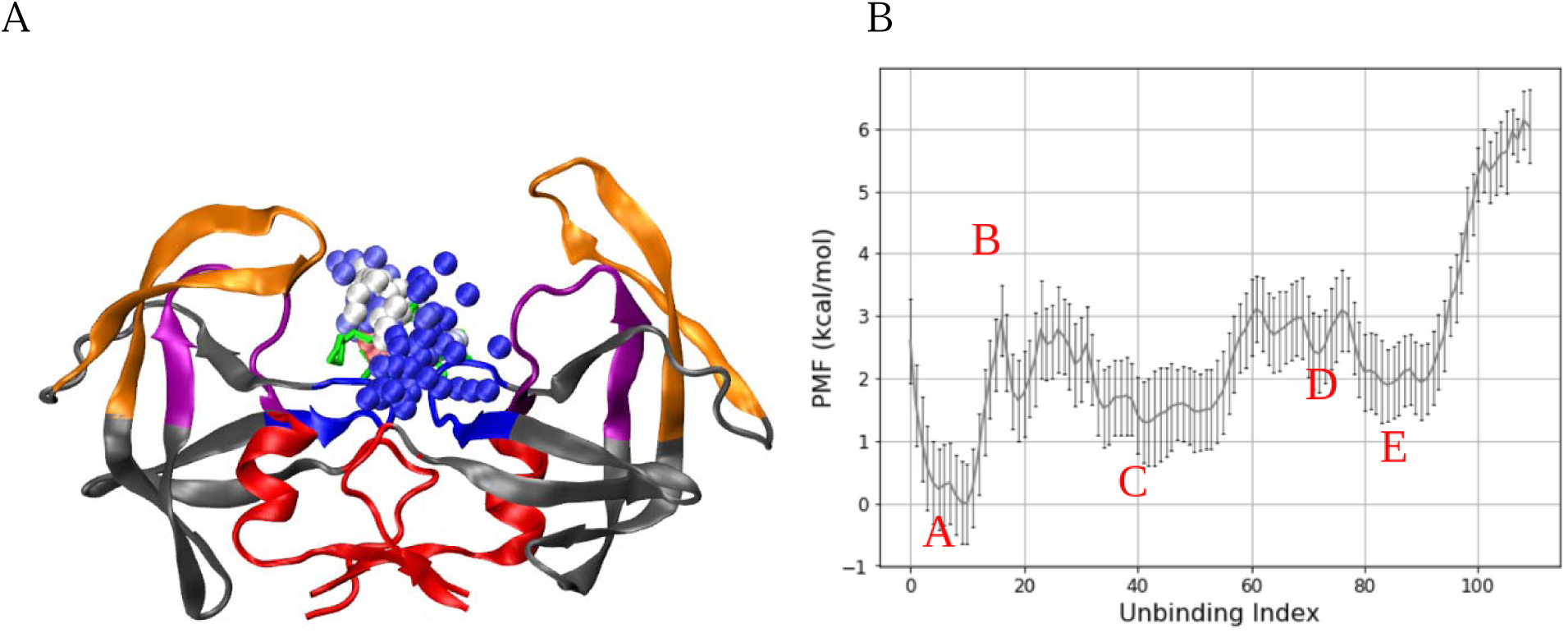
Free energy landscape of ritonavir unbinding from HIVp under pathway C. **(A)** Ritonavir dissociation under pathway C. HIVp colored as in Figure 1. Each bead represents a position of center of mass of ritonavir with 1-ns interval taken from aMD dissociation, with total simulation time of 543.1 ns. The HIVp conformation is taken from the final frame of the dissociation trajectory. Color beads present center of mass of ritonavir taken from frames in the beginning (red), middle (white) and near the end (blue) of the trajectory. Ritonavir’s initial position is green licorice. **(B)** Free energy landscape of ritonavir under pathway B with labeled maxima and minima. Major energy barriers (B) and local minimum (A,C,D,E) labeled in red, and the conformations are detailed in **Figure 8**.

**Figure 8.**
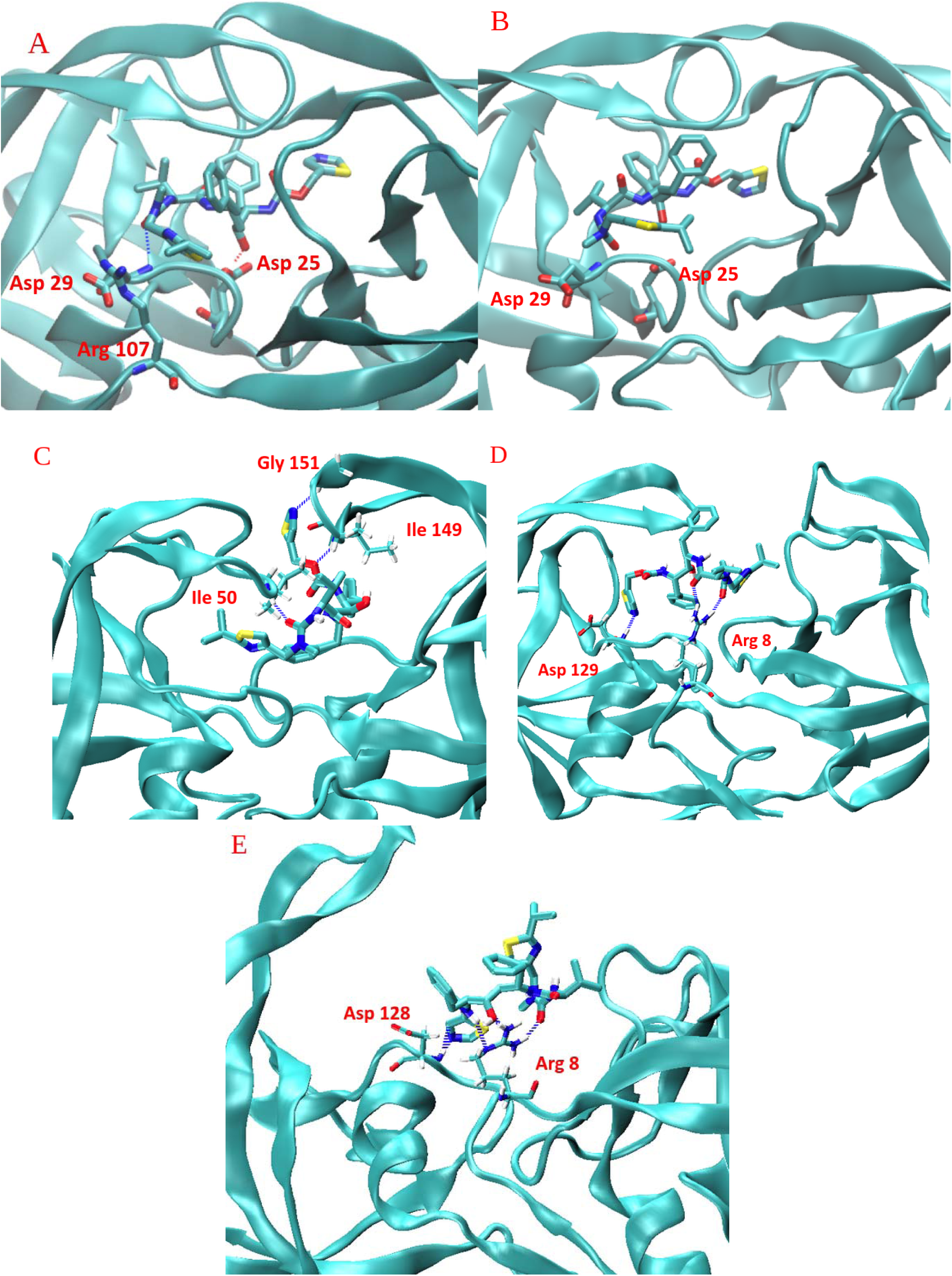
Progression of molecular interactions of Ritonavir and HIVp during dissociation via pathway. **C.** Local energy minima or barriers are indicated in **Figure 7**. **(A, C, D, E)** Local free energy minima are explained via bound-state stabilization by Hydrogen bonds between ritonavir and HIVp. **(B)** Free energy barrier can be described by the breaking of favorable interactions.

#### Insights provided from free energy plot

Energy barriers and wells were successfully attributed to particular protein-ligand interactions, validating the free energy plot specific to each dissociation pathway. These interactions contribute highly to the overall unbinding free energy. Hydrogen bonding was seen to have a large impact of movement in the flap region. However, visual analysis of protein-ligand interactions did not account for the discrepancy in occurrence between Pathway A and the other two Pathways. It is likely that an additional component, separate from unbinding energy, is impacting the dissociation of ritonavir from HIVp.

#### Analysis of pathway-specific side-chain correlation

Mutation studies have shown that single mutations are not impactful on HIVp affinity for substrate, however mutation combinations result in drug resistance (43). This indicates that single residue studies are not sufficient on their own in understanding the ways in which ritonavir can dissociate from HIVp. In combination with the previous single residue studies, we ran side-chain correlation analysis specific to each of the three pathways. This gives insight into the cohesiveness of the protein in binding ritonavir. Pathway A exhibited an extensive network of highly correlated residues (**Table 2**).

**Table 2.** Residues with substantial correlation in Ritonavir-HIVp dissociation Pathway A. Positive correlation cutoff > 0.50. Negative correlation cutoff < -0.50. 10 residue pairs had significant positive correlation. 8 residue pairs had a significant negative correlation.

| Positive Correlation (>0.5) |  |  | Negative Correlation (<-0.5) |  |  |
| --- | --- | --- | --- | --- | --- |
| Residue | Residue | Correlation | Residue | Residue | Correlation |
| ILE66 | ASN83 | 0.601541 | TYR59 | ASN83 | -0.52429 |
| VAL75 | GLN160 | 0.529861 | ILE62 | TRP141 | -0.51558 |
| VAL82 | GLN101 | 0.501499 | ILE62 | LEU122 | -0.599 |
| ASN83 | ASN182 | 0.573296 | ASN83 | ASN182 | -0.5665 |
| ASN83 | ILE161 | 0.530882 | ASN83 | ASN182 | -0.6823 |
| ASN83 | ASN182 | 0.644101 | ILE84 | ILE165 | -0.50422 |
| ARG87 | ILE112 | 0.526643 | GLN92 | ILE183 | -0.51592 |
| ILE114 | MET135 | 0.508678 | ILE112 | ASN83 | -0.56379 |
| LEU122 | ASN182 | 0.547785 |  |  |  |
| VAL131 | ILE171 | 0.505891 |  |  |  |

These residues had high coverage across the majority protein and were not localized to any singular region, though they were primarily absent in flap regions (**Figure 9**). Higher amounts of correlated residues were found on the side of HIVp in which ritonavir dissociates along in pathway A. Correlation was demonstrated in mirror loop residues ASN83 and ASN182, indicating symmetric loop movement during ritonavir dissociation. Pathway A demonstrates a highly correlated protein that arranges side-chain movements to interact with ritonavir. The flap regions move according to single residue conflicts and are predominantly governed by hydrogen bonding as it relates to the PMF. However, in Pathway A, the rest of the protein exhibits organized motion that helps to shape HIVp and stabilize transient states of ritonavir. This correlation network cannot be predicted using free energy alone. These findings are supported by known resistance data, as residues appearing in the correlation network, ILE84 and Val131, are known to have less favorable interactions in drug resistant strains of HIVp (43). It is worth noting that when the correlation threshold is lowered below |0.5|, this correlation network is far more expansive and encompasses more known resistance residues.

**Figure 9.**
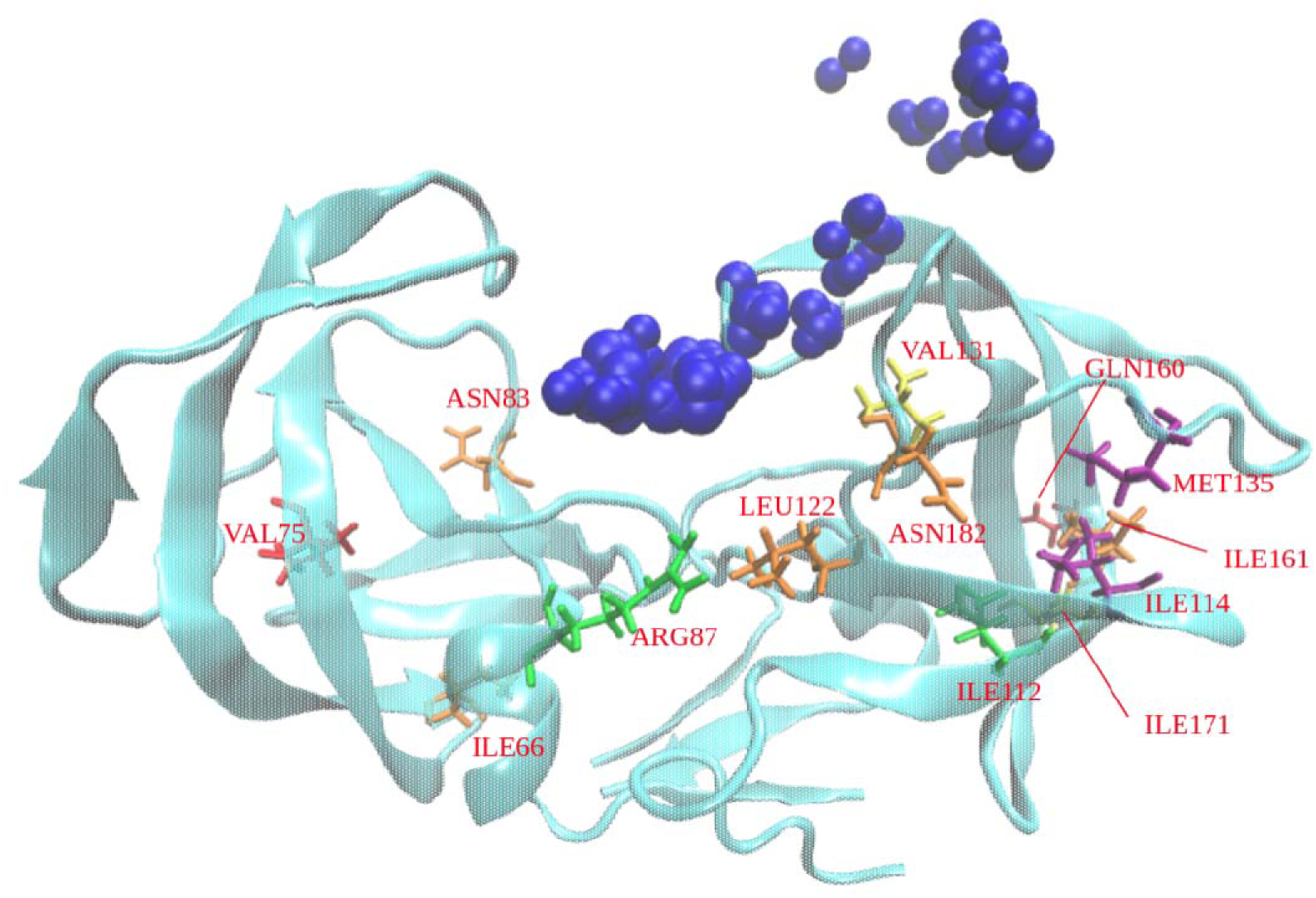
Highly positively-correlated residue in Ritonavir-HIVp dissociation Pathway. **A.** HIVp shown in cyan. Ritonavir position during dissociation taken from representative frames shown in bead illustration (grey), dissociating between the flap and the loop regions. Crude approximate binding spheres are used to show general ritonavir location. Image captures backbone and sidechain positions at the final unbinding index. Residues are dispersed in all regions excluding the flap regions. Increased number of correlated residues on side in which ritonavir dissociates. Sidechain colors reflect correlation pairings/groups. Pathway B exhibited a less extensive network of highly correlated residues (Table 3). These residues were localized primarily to the interface region and once again were absent in flap regions (**Figure 10**). Pathway C lacked a correlation network. Only two residue pairs in pathway C demonstrated a high positive correlation. Pathway c was absent of negatively correlated residues (Table 3).

**Table 3.** Residues with substantial correlation in Ritonavir-HIVp dissociation pathway B and pathway C. Positive correlation cutoff >.50. Negative correlation cutoff <-.50. Pathway B shows 3 residue pairs with significant positive correlation and 1 residue pair with a significant negative correlation. In Pathway C, 2 residue pairs were identified with a significant positive correlation and 0 residues shared a significant negative correlation.

| Pathway | Residue | Residue | Correlation |
| --- | --- | --- | --- |
| B | THR31 | LEU89 | 0.523319 |
| B | LEU89 | ILE184 | 0.625443 |
| B | ILE184 | LEU188 | 0.654553 |
| B | ILE183 | LEU189 | -0.50999 |
| C | LEU19 | LEU23 | 0.504406 |
| C | ASP30 | THR91 | 0.506992 |

These residues did not follow any clear pattern or localization and appear random in nature (**Figure 11**). Neither pathway B nor C had a correlation network involving any residues known to impact drug resistance. Single residues give some insight into the total energy of each dissociation pathway; however, the exceptionally cohesive motion of the protease seems to be the major contributor to Pathway A being favored.

**Figure 10.**
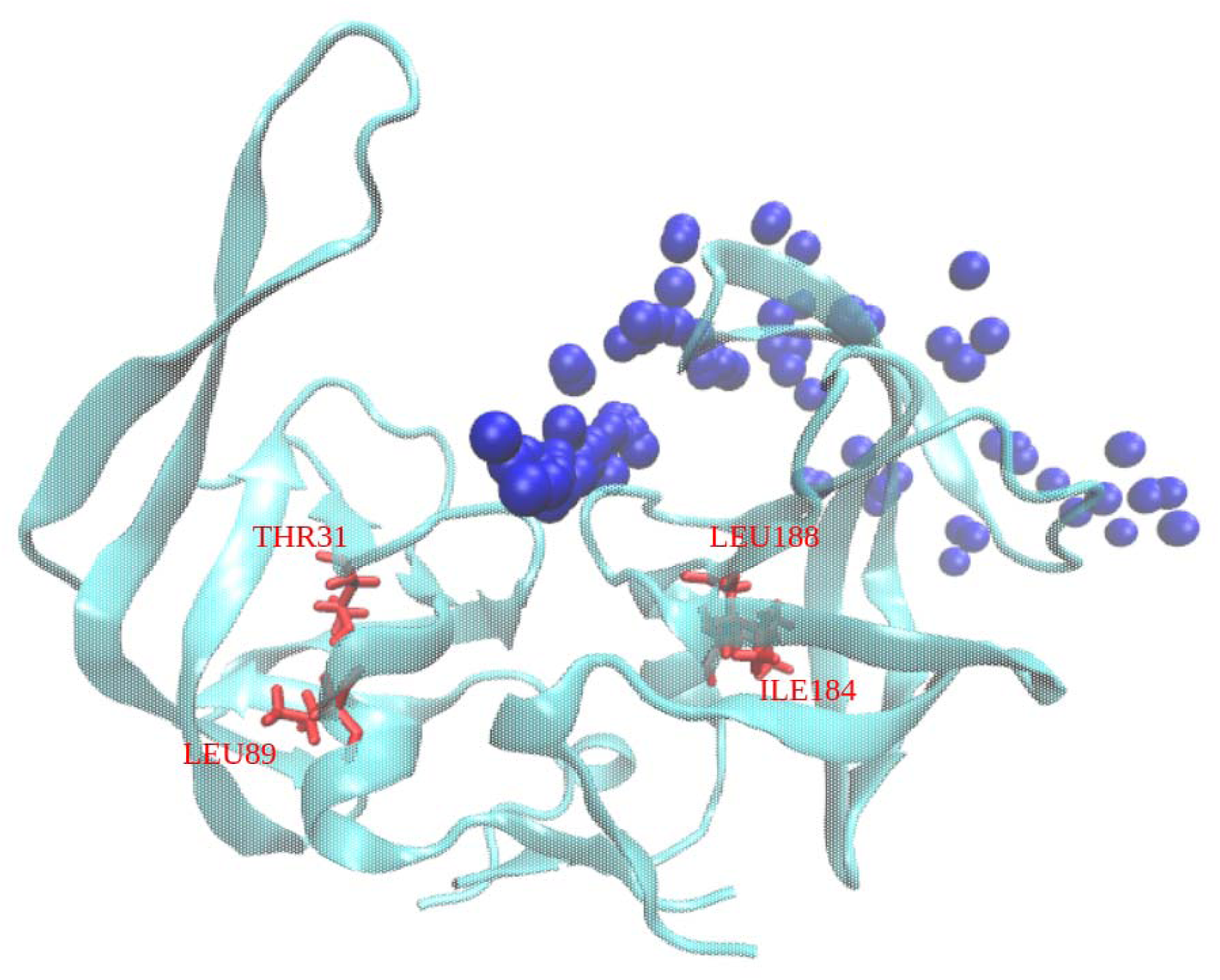
Highly positively-correlated residue in Ritonavir-HIVp dissociation Pathway. **B.** HIVp shown in cyan. Ritonavir position during dissociation taken from representative frames shown in bead illustration (grey), dissociating along the flap region. Crude approximate binding spheres are used to show general ritonavir location. Image captures backbone and sidechain positions at the final unbinding index. Correlated residues are primarily localized to the interface region and are absent from the flap region. Only one correlation group is present (red).

**Figure 11.**
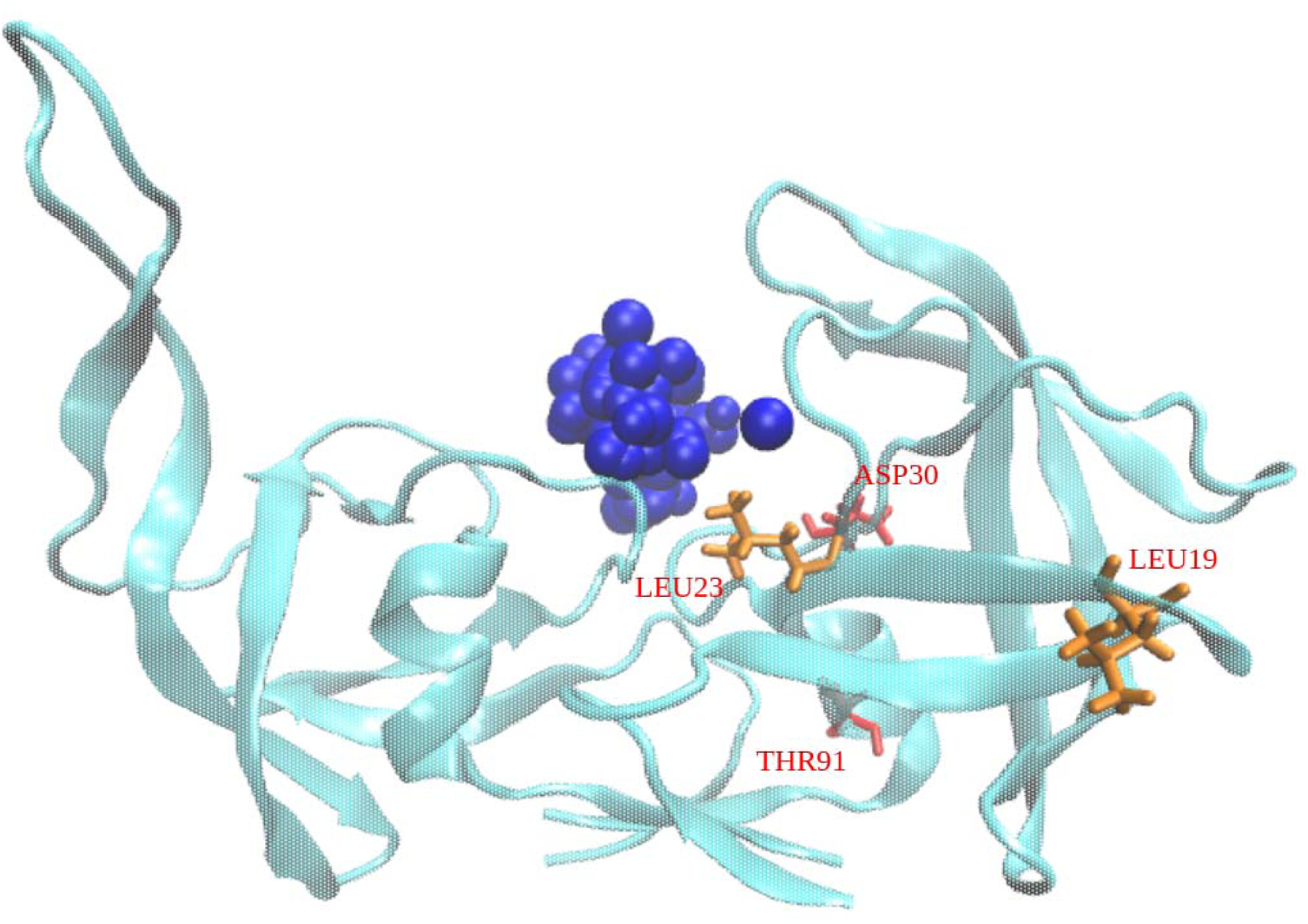
Highly positively-correlated residue in Ritonavir-HIVp dissociation Pathway. **C.** HIVp shown in cyan. Ritonavir position during dissociation taken from representative frames shown in bead illustration (grey), dissociating on the interface region. Crude approximate binding spheres are used to show general ritonavir location. Image captures backbone and sidechain positions at the final unbinding index. Sidechain colors reflect correlation pairings/groups. Correlated residues show no distinct localization.

## CONCLUSIONS

Due to increasing drug resistance from protein mutations, it is increasingly challenging to develop drugs with high efficacy when a protein target mutates. Considering drug binding kinetics by examining the drug binding/unbinding pathways reveals transient states during drug binding/unbinding, which provide a more complete picture of molecular recognition and understanding into mutation effects of distal sites. While many protein systems demonstrate a single channel for ligand binding/unbinding, HIVp has a highly flexible flap regions to allow ligand binding/unbinding via different pathways. As a result, examining one single-pathway is insufficient to understand drug unbinding. This study therefore investigated different unbinding pathways in detail to uncover key transient states and residues of interest. By combining this with sidechain correlation, we uncovered new information on the motion of HIVp as ritonavir dissociates. Our study shows that the computed free energy landscape captures detailed ligand-protein interactions during ligand unbinding using 2D PC space to define unbinding coordinates and milestones. However, the computed absolute binding free energy and residence time were at moderate accuracy. We also used ligand RMSD as unbinding coordinates, and use of the 1D milestones resulted in accurate absolute binding free energy and drug residence time, compared with experimental measurements. Combining these two approaches, we validated our sampled unbinding pathways and demonstrated key interactions that brought free energy barriers and minima during ritonavir dissociation.

Ritonavir-HIVp interactions, particularly those in the flap region, show that hydrogen bonding contributes significantly to free energy and governs dissociation of ritonavir from HIVp. These interactions can be directly tied to the unbinding free energy plot in each of the three pathways. However, the studies of ritonavir-HIVp interactions are insufficient to explain the low occurrence of Pathway B and Pathway C in comparison to Pathway A, considering the similarity in unbinding free energy between the pathways. We therefore carried out side-chain correlation to find residue-residue interaction network, which is not directly visible in the energy profile. Pathway A exhibits remarkably large residue correlation in the protease itself, allowing HIVp to adjust intramolecular interactions to better accommodate ritonavir to form hydrogen bonds during ligand dissociation and lead to the most popular unbinding pathway. The correlation analysis disclosed these highly correlated residues in Pathway A, which include ILE84 and Val131 reported in mutation and drug resistance. When the residues mutated, the residue correlation network was altered, leading to less stable intermediate states that accelerates drug dissociation. This study explains how the distal sites may affect drug binding affinity. As the protein most commonly works via a widescale correlation network, we also demonstrated that considering single residue-drug contacts is not enough in understanding drug unbinding processes. This explains why multiple mutations need to happen simultaneously to cause drug resistance to ritonavir. Our study also explains when mutations occur outside of flap regions, though not directly affecting drug-HIVp binding in the catalytic pocket, it may disrupt the correlation network of HIVp and facilitate the dissociation of ritonavir. These mutation effects would be difficult to predict using thermodynamic data alone. Based on the results of this study, protein correlation may be used as a factor in predicting ligand-protein binding kinetics and specificity. If a ligand-protein system has a widespread correlation network, it may suggest that it can better stabilize transient states, thus making the ligand an ideal inhibitor with high residence time.

## DATA AVAILABILITY

All computational data and input files are available upon request.

## ACKNOWLEDGMENTS

This study was supported by the US National Institutes of Health (GM-109045). Computations were performed using the computer clusters of the University of California, Riverside High Performance Computer Cluster and NSF national supercomputer centers ACCESS (CHE130009).

