## Supplementary Material for "Pathway Specific Unbinding Free Energy Profiles of Ritonavir Dissociation From HIV-1 Protease"

| | Initial Frames taken every 100 ps | Inital Frames taken every 20 ps |
|---|---|---|
| Pathway | Simulation time (ns) | Simulation time (ns) |
| A | 0 to 200 | 200.02 to 248.50 |
| B | 0 to 275 | 275.02 to 304.90 |
| C | 0 to 500 | 500.02 to 543.10 |

**Table S1. Frequency of initial frames specific to each dissociation pathway.** Frames are taken less frequently during the beginning of each run. To better sample PC space, the interval of initial frame selection is decreased to 20ps once the ligand begins to leave the pocket. Initial frames varied between pathways to account for differences in simulation length and ligand residence time.

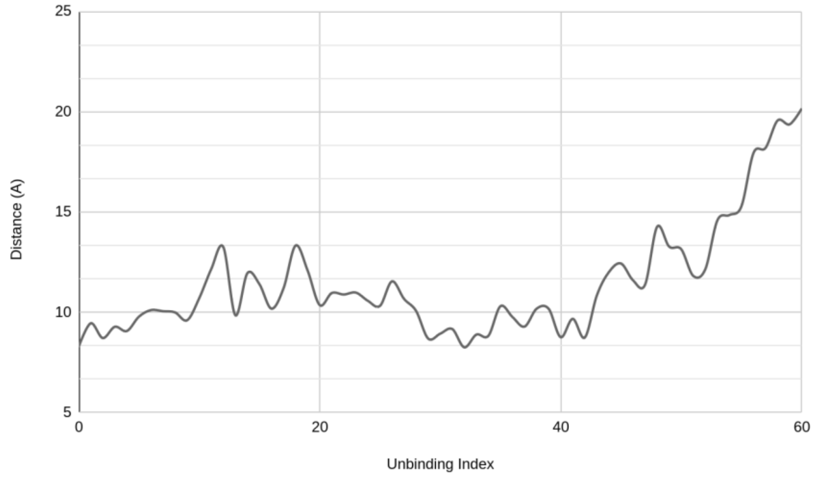


**Figure S1: Distance between ritonavir and HIVp during unbinding pathway A**
Distance between HIVp and ritonavir is measured for the first 60 unbinding indexes out of 91 total representative frames. This is used to capture approximate distance in the bound states in addition to important transient states. Distance is calculated using the approximate center of mass. HIVp center of mass is estimated as the alpha carbon of GLY27. Ritonavir center of mass is estimated using C15. Distance between ligand and protein consistently rises and falls, indicating that ritonavir moves back and forth within the pocket.

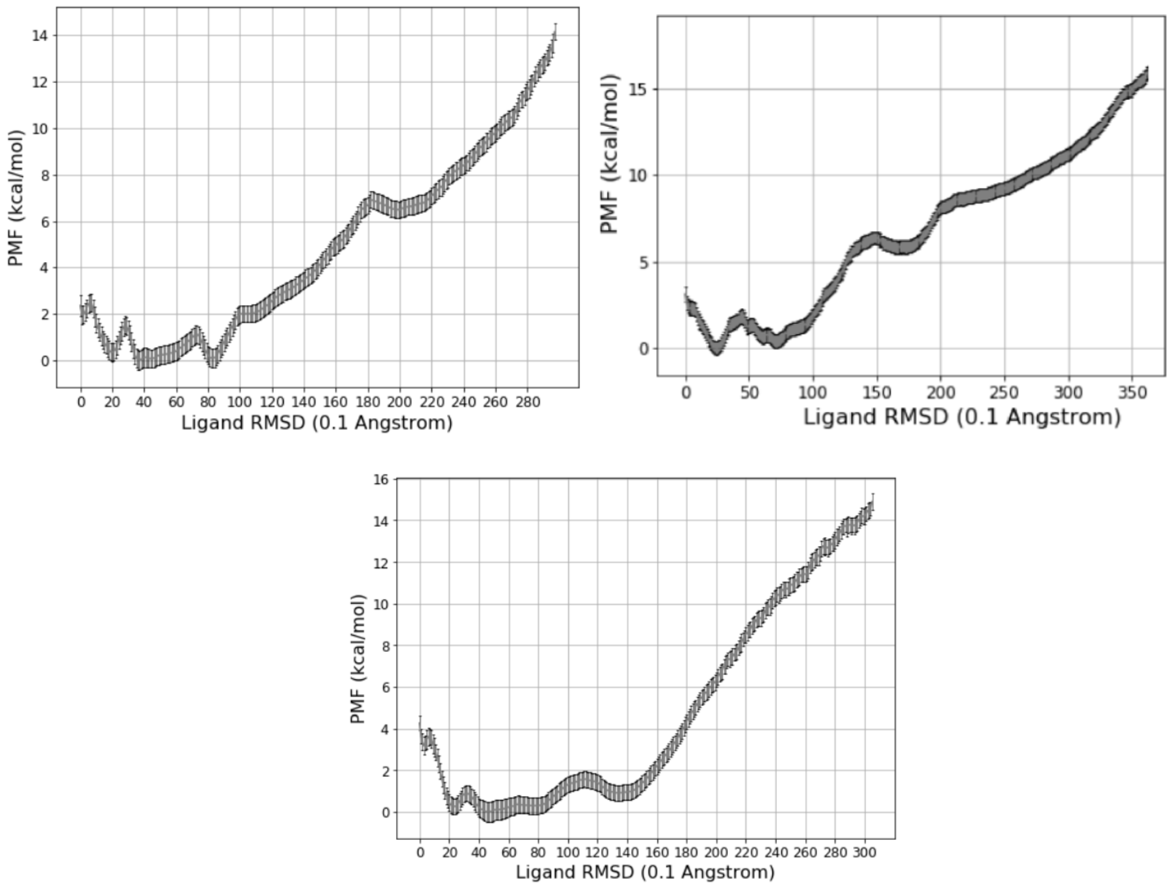


**Figure S2. Pathway specific free energy landscape of ritonavir unbinding from HIVp.** Free energy landscape of pathway A (top left), pathway B (top right), and pathway C (bottom), derived from RMSD method. Energy values derived using this method agree with experimental findings, validating the dissociation trajectory used.

A

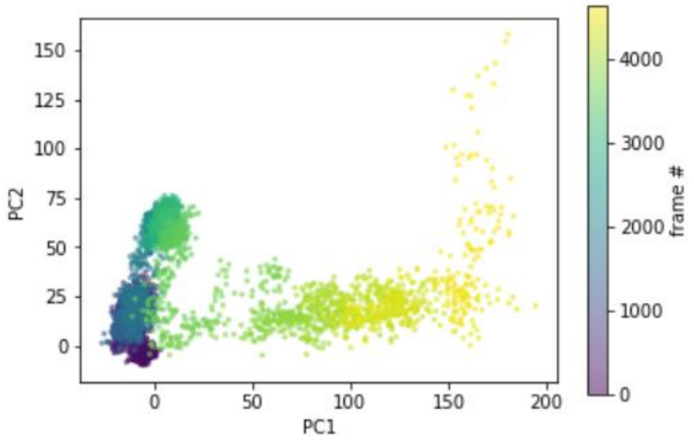


B

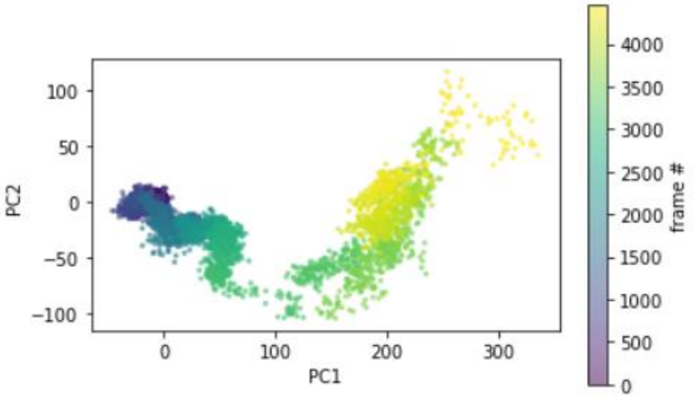


C

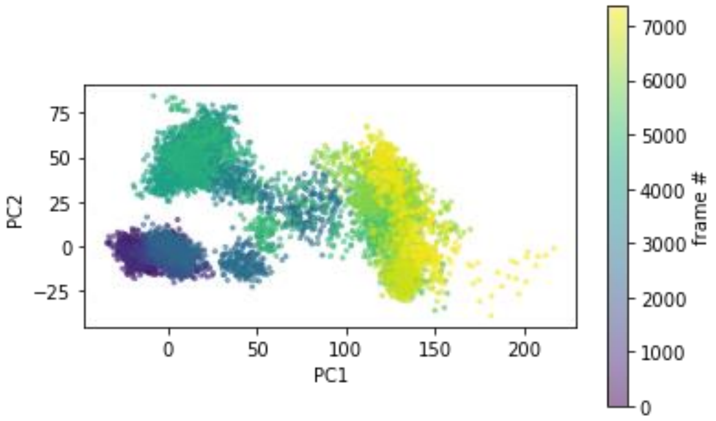


**Figure S3. Projection of PC1 and PC2 onto PC space.** PC visualization shown for pathway A (top), pathway b (middle), and pathway c (bottom). Due to high coverage calculated by the explained variance ratio, only the first two PCs are necessary to capture most key movements during ritonavir dissociation from HIVp.

A

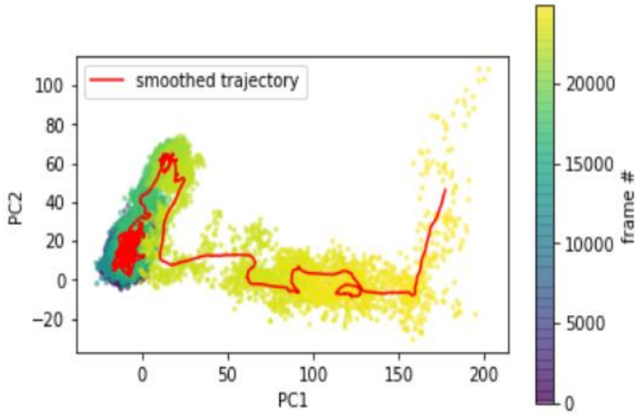


B

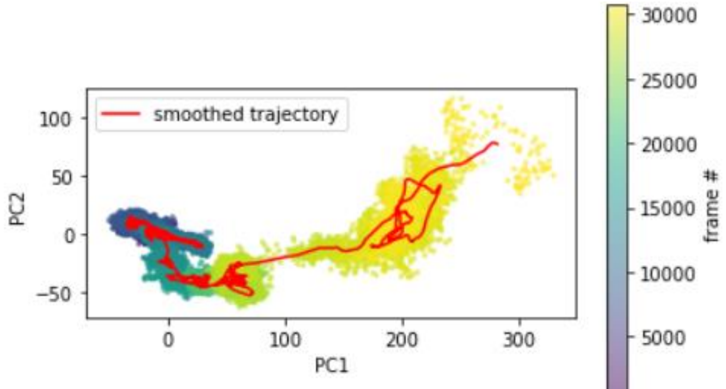


C

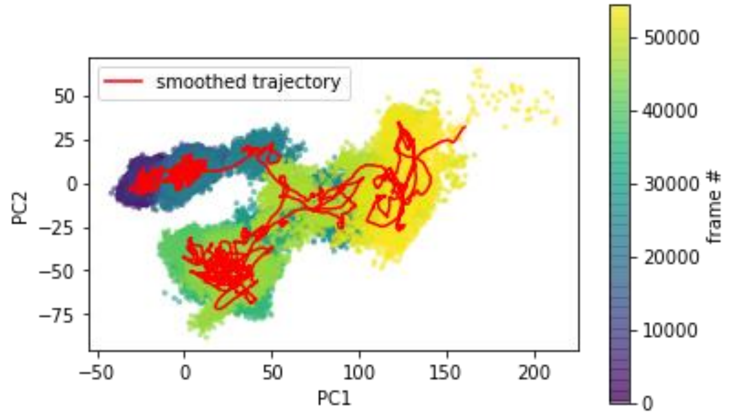


**Figure S4. Projection of smoothed dissociation trajectory onto PC space.** Smoothed dissociation trajectory shown in red. PC visualization shown for pathway A (top), pathway b (middle), and pathway c (bottom). We use low-d projections to sketch a reaction coordinate. A smooth path is generated by averaging frames in the before and after 100 steps of each frame

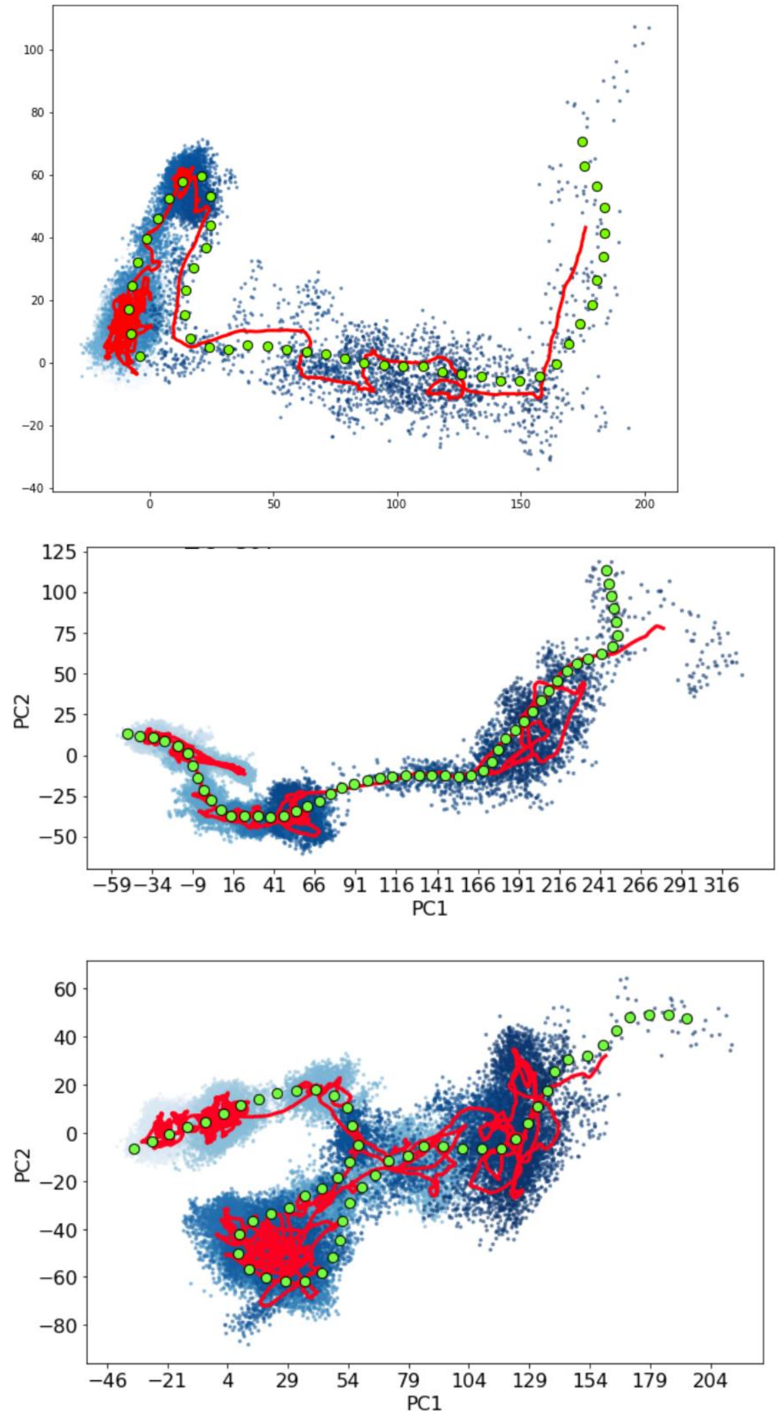


**Figure S5. Final reaction path for pathway A,B, and C.** Final path shown for pathway A (top), pathway b (middle), and pathway c (bottom). Smoothed dissociation trajectory shown in red is generated by averaging frames in the before and after 100 steps of each frame. Center of milestones (green) is set at a width of 4 angstrom.

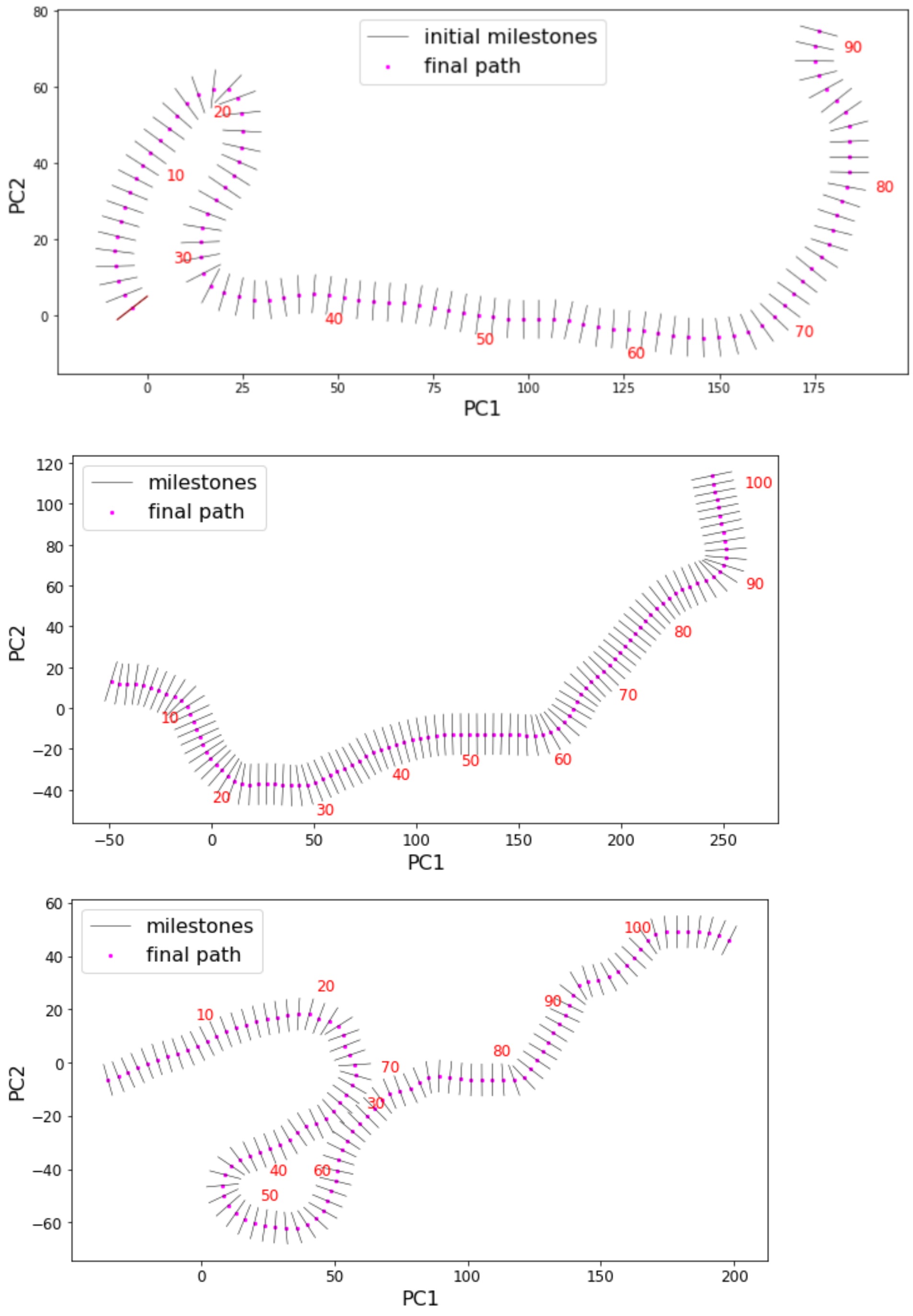


**Figure S6. Optimized milestones shown on PC space.** Milestones shown for pathway A (top), pathway b (middle), and pathway c (bottom). We perform both global and local optimization of the slopes of milestones to 1) avoid crossing of milestone and 2) to make sure all the milestone cells are trapezoid.

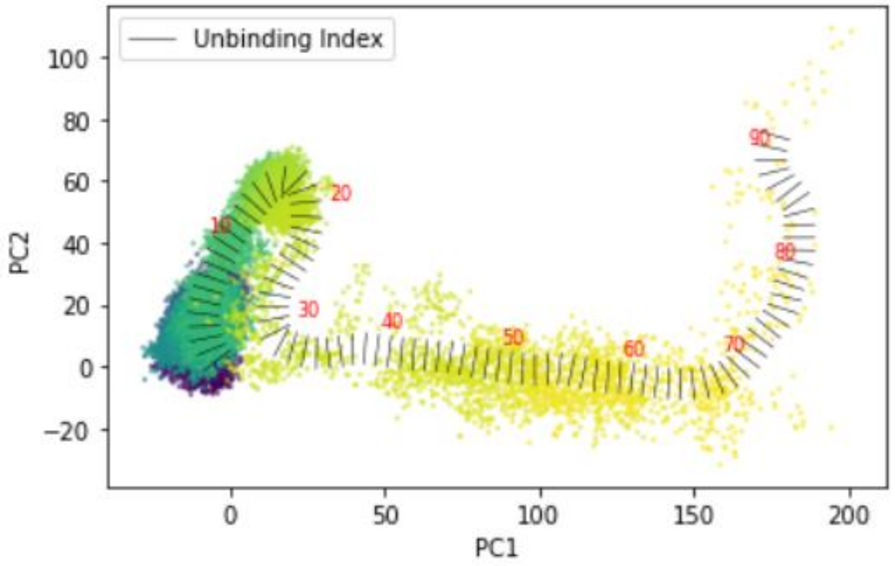


**Figure S7. Optimized milestones shown over PC projection.** Sample visualization of Pathway A. Milestones (figure S6) shown over PC projection (figure S3)

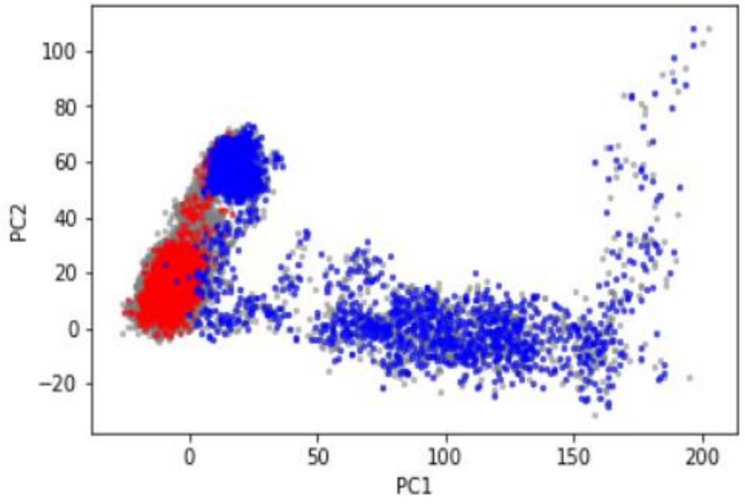


**Figure S8. Initial frames taken to run short MD.** Initial frames taken to run short MD for pathway A. Frames at the beginning of the simulation are taken every 100ps (red). Frames occuring once ritonavir begins to dissociate are taken evet 20ps (blue).

A

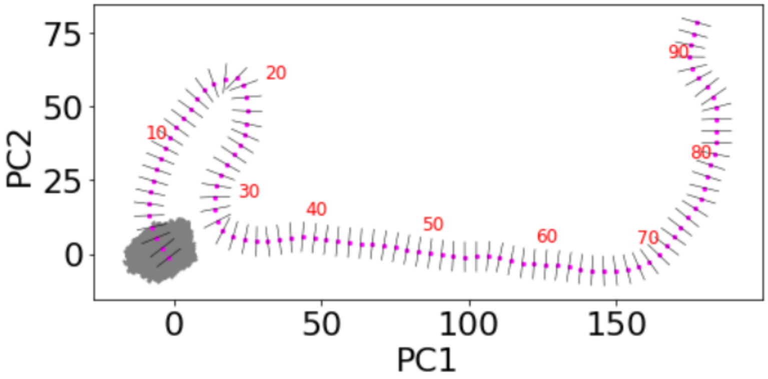


B

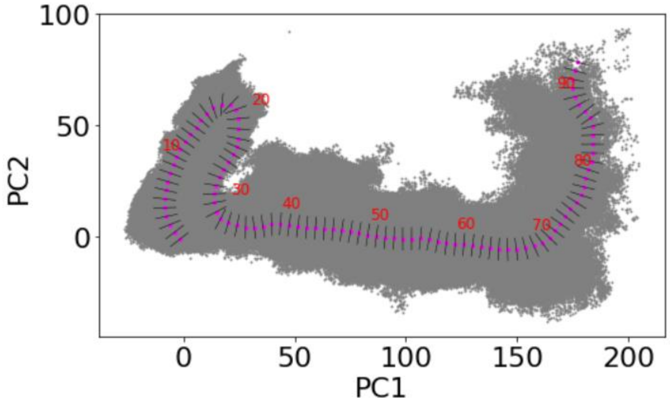


**Figure S9. Projection of short MD onto PC space.** Short MD from pathway A are projected onto the respective PC space. **(A)** First short MD projected onto PC space. **(B)** All short MD projected onto PC space.

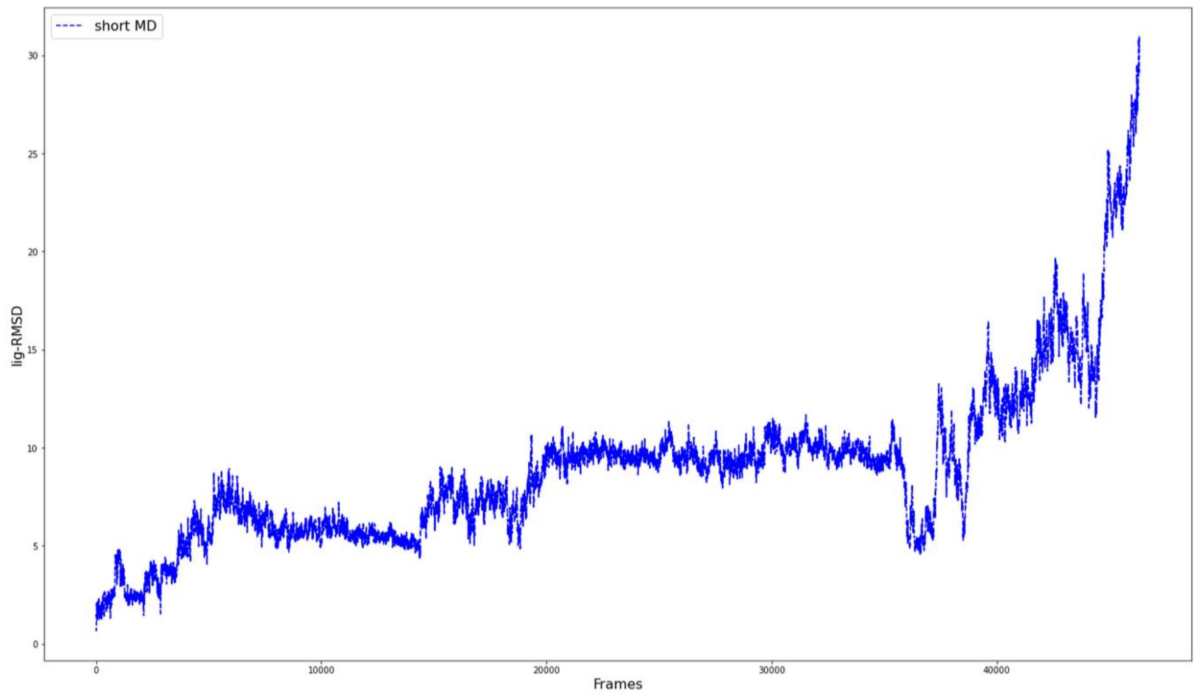


**Figure S10. Ritonavir RMSD during Pathway A.** Ritonavir RMSD calculated for pathway A using short MD. This is used to calculate an RMSD based free energy plot.
